# A conserved logic for the development of cortical layering in tetrapods

**DOI:** 10.1101/2025.10.01.679862

**Authors:** Astrid Deryckere, Saket Choudhary, Connor Lynch, Lian Kirit V. P. Limperis, Pauline Affatato, Jamie Woych, Elias Gumnit, Alonso Ortega Gurrola, Rahul Satija, Christian Mayer, Maria Antonietta Tosches

**Author notes:** Laboratory of Reproductive Genomics, KU Leuven; Leuven, 3000, Belgium. Koita Centre for Digital Health, IIT Bombay; Mumbai, 400076, India.

## Abstract

The cerebral cortex is part of the pallium, a brain region conserved across vertebrates yet remarkably diverse in structure and cellular composition. A defining feature of the cerebral cortex is its organization into neuronal layers with distinct gene expression profiles, input-output connectivity, and function. According to prevailing models, the cerebral cortex emerged in ancestral amniotes (mammals and reptiles) following innovations in pallial development that enabled the generation of diverse neuron types and their laminar organization.^1–11^ However, little is known about pallial development and architecture in amphibians, the sister group of amniotes. Here we show that in the salamander *Pleurodeles waltl*, the dorsal pallium is organized in distinct superficial and deep layers with neurons that develop following cellular and molecular principles of mammalian corticogenesis. Using birthdating analysis, barcode-based lineage tracing, and single-cell RNA sequencing, we find that radial glia temporal states and intermediate progenitor cells are conserved across species, while neuronal differentiation trajectories are highly evolvable. Neurons generated at different developmental time points occupy different layers and exhibit distinct molecular and projection identities. Thus, temporally-patterned neurogenesis represents an ancient organizing principle of layered pallia, although mammals display an inverted layer order along the radial axis. Together, these findings demonstrate that the core developmental principles underlying cortical layering - including temporal patterning, intermediate progenitors, and laminar organization - predate the origin of amniotes. Our results suggest that the evolutionary expansion of the mammalian neocortex built upon a deeply conserved developmental framework already present in early tetrapods.

## Introduction

The laminar organization of the cerebral cortex is a defining feature of mammalian brains, underpinning complex information processing and higher cognitive functions. Exactly how the cortex arose and diversified during vertebrate evolution remain long-standing questions in neuroscience. The cerebral cortex is part of the pallium, the dorsal subdivision of the telencephalon. The pallium exists in all vertebrates but its anatomical organization is extremely diverse across species.^11^ While some regions of the pallium do have neurons arranged into nuclei, the cerebral cortex is anatomically defined as “*a neural aggregate that consists of alternating layers of different neuronal populations and their associated fibers*”.^12^ According to this established definition, the only vertebrates with a true cerebral cortex are mammals and non-avian reptiles (lizards, turtles, crocodiles) though a layer-like organization may have evolved independently in birds.^13–15^ The cerebral cortex has thus long been considered an amniote innovation, absent in anamniotes such as amphibians.

Cortical layers are not just an anatomical feature; they are part of the implementation of a successful biological architecture for associative learning.^16–19^ In all cortices, fibers are arranged in an orthogonal grid, in which the pyramidal neurons’ dendrites extend along the radial axis (from ventricle to surface) and the incoming axons extend along the tangential axis (parallel to the cortical surface). Inputs from distinct brain regions target specific cortical depths, where they synapse on the dendrites of broadly distributed neuronal populations. As a result, neurons in different layers integrate distinct sets of inputs, and each neuronal layer functions as a parallel processing unit.^20^ This architecture makes the cerebral cortex an efficient, flexible, and expandable layout for associative learning and memory storage.^21–24^ Understanding the origin and evolution of cortical layering is then essential for elucidating the emergence of cortical computations.

Reflecting their functional specialization, cortical layers are made of glutamatergic neuron types with distinct morphologies, gene expression, and input-output connectivity. The presence of multiple glutamatergic types characterizes not only the neocortex, with its six layers, but also simpler cortices in mammals and non-avian reptiles. In addition to the neocortex, mammals also have cortical regions with fewer layers,^25^ such as the hippocampus (“archicortex”) and the piriform cortex (“paleocortex”). These simple mammalian cortices anatomically resemble the cerebral cortex of non-avian reptiles, where a cellular layer is separated from the ventricular and the pial surfaces by fibers (axons and dendrites).^16,26–28^ Detailed anatomical and molecular studies, however, have shown that even these simple reptilian and mammalian cortices comprise sublayers of glutamatergic neurons with distinct gene expression profiles, input-output connectivity, and functions.^15,28–35^ Therefore, the presence of distinct types of glutamatergic neurons organized into layers is a conserved feature of the cerebral cortex in amniotes.

In contrast, layers of distinct neuron types have not been described so far in the pallia of fish and amphibians (anamniotes). In teleost fishes such as zebrafish and goldfish, the pallium is an aggregate of nuclei, while in lampreys, certain sharks, lungfishes, and amphibians, it comprises densely-packed cell bodies near the ventricle and a fiber-rich zone closer to the pial surface.^11,36–38^ The observation that the pallium is not layered in anamniotes has led to the conclusion that a layered cerebral cortex evolved only in amniote ancestors.

The layered architecture of the cerebral cortex results from the mechanisms that generate neuronal diversity during development. For this reason, it is believed that the cerebral cortex arose in evolution through innovations of developmental programs in the pallium that enabled the generation of multiple types of neurons, and thus, layers.^1–11^ This hypothesis, however, remains largely untested because pallial neurogenesis has been poorly studied in fish and amphibians. During cortical development, the generation of multiple neuron types and their layered arrangement are based on three principles: (1) the generation of distinct neuron types in a stereotyped temporal sequence by multipotent radial glia progenitors (temporal patterning);^39–41^ (2) the amplification of the neuronal output of radial glia by intermediate progenitor cells (IPCs);^8,40^ (3) the final distribution of distinct neuron types at different distances from the ventricle, either through “outside-in” neurogenesis (with early-born neurons farthest away from the ventricle, as seen in the reptilian dorsal cortex)^29,42,43^ or radial migration and “inside-out” neurogenesis (with early-born neurons closest to the ventricle, as seen in most mammalian cortices)^44,45^. As a result, neuronal birthdate is tightly linked to molecular identity, projection pattern, and layer allocation.^27,44,46^

Data from amphibians and fish indicate outside-in neurogenesis in the pallium,^47–49^ but the presence of multipotent progenitors and neuronal diversity remain largely untested in anamniotes, and the evolutionary history of IPCs controversial.^3,5,7,9,50,51^ Here, we characterize pallial development in an amphibian, the salamander *Pleurodeles waltl*. Our cellular and molecular analysis shows the presence of multipotent radial glia progenitors and IPCs, the sequential generation of two neuron types with distinct transcriptomes and projections, and the arrangement of these neuron types into two layers. Furthermore, we show that salamander and mouse radial glia express a common set of temporally-modulated genes, and that early- and late-born neurons share not only gene module expression but also projection patterns across species. Our analysis suggests that, despite their transcriptomic divergence, pallial neuron types in amphibians, reptiles, and mammals develop according to the same logic, and that their birthdates, and not absolute position along the radial axis, reflect common evolutionary origins. Our findings demonstrate that multipotent radial glia, temporal patterning, IPCs, and layering are present in the amphibian pallium, challenging the notion that the cerebral cortex is an amniote innovation, and uncovering the developmental changes that likely enabled the exceptional expansion and diversification of the mammalian neocortex.

## Results

### The adult salamander dorsal pallium includes two layers of glutamatergic neuron*s*

Recently, we characterized the transcriptomes and spatial distribution of neuron types in the *Pleurodeles* telencephalon, and discovered that their pallium is organized in molecularly-distinct regions along the medio-lateral axis.^52^ Here we focus on the dorsal pallium (DP), the pallial region nested between the medial pallium (hippocampal homolog) and the olfactory pallium (lateral and ventral pallium), and thus analogous, for its position, to the reptilian dorsal cortex and mammalian neocortex. The salamander dorsal pallium is not homogeneous, but includes six distinct clusters of glutamatergic neurons: telencephalic glutamatergic (TEGLU) neuron clusters 8-11, 13 and 20.^52,53^ Here, we further analyzed the spatial distribution and differential gene expression of these DP clusters, and found that they could be grouped in superficial layer (DP-SL) and deep layer (DP-DL) types, based on the differential expression of hundreds of genes, including transcription factors, neuropeptides, and proteins involved in axonal pathfinding and synaptogenesis (**Fig. 1A–F**, **fig. S1A-C**). Superficial-layer neurons (cluster TEGLU20) uniquely expressed the neuropeptide precursor gene *Nts* and the transcription factor *Foxp1* at high levels, and were transcriptomically distinct from other *Nts*-expressing types in adjacent pallial regions (**fig. S1C-F**) and from sparse *Nts*-expressing GABAergic interneurons in deep layers (**Fig. 1E’’**). Conversely, deep-layer neurons (clusters TEGLU 8-11, 13) uniquely expressed protein phosphatase and calcium sensing proteins *Ppm1h* and *Otof* (**fig. S1C**). These neurons showed further heterogeneity based on their anterior-posterior and mediolateral position. For our developmental analysis, we focused on the anterior DP, where *Nts*-expressing neurons are more abundant. The anterior DP was identified by the expression of *Frmd8* in the deep layer; these neurons also expressed the transcription factor *Nfia* (clusters TEGLU8, 10, 11, **Fig. 1D-F**, **fig. S1G**). The posterior DP, demarcated by *Nr2f2* expression (cluster TEGLU9), and a population of *Calb2* expressing neurons (cluster TEGLU13) at the boundary between DP and lateral pallium (LP) (**fig. S1C** and Woych et al.^52^), were excluded from subsequent analysis.

**Fig. 1.**
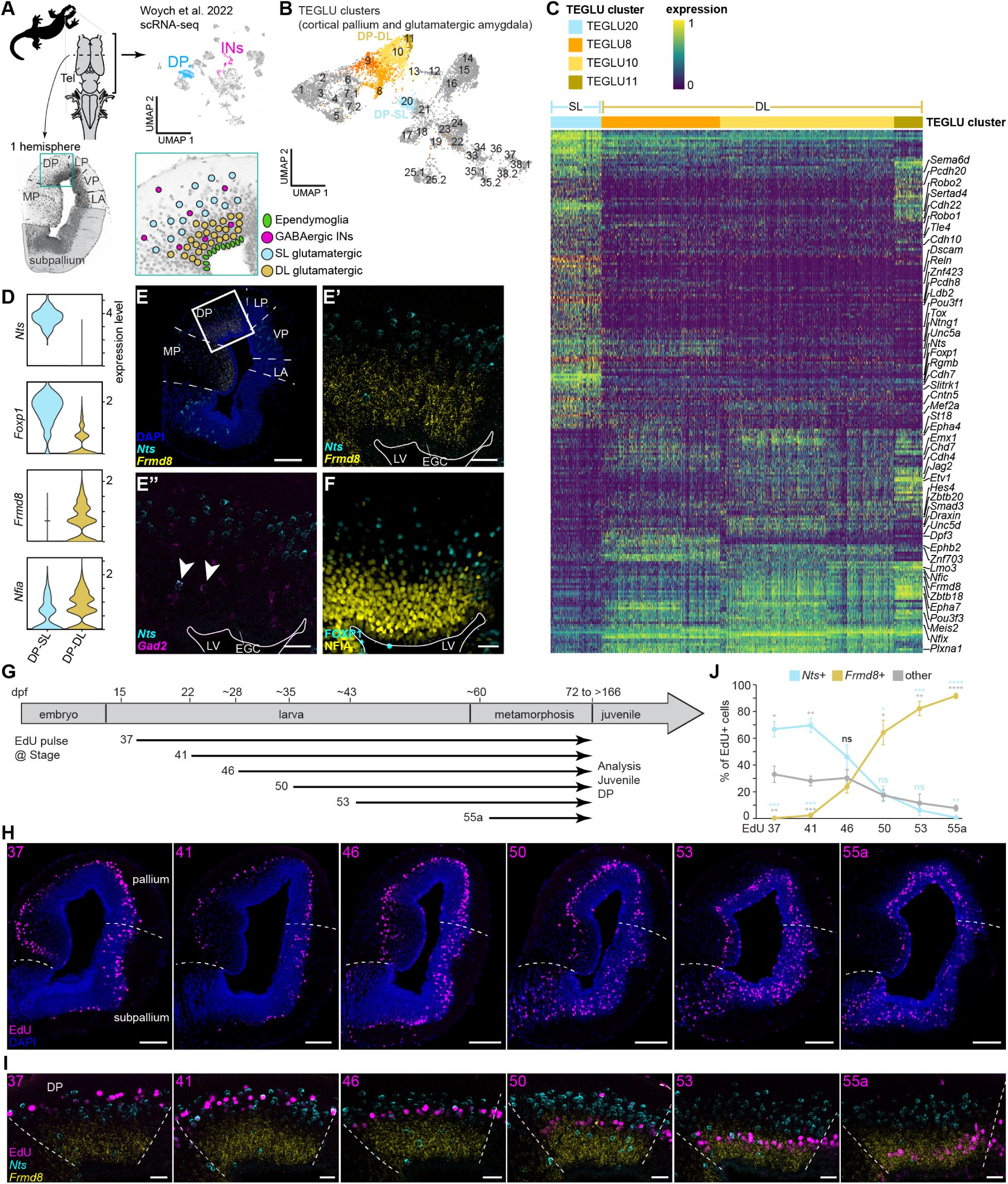
Two distinct layers of glutamatergic neurons are generated sequentially in the salamander dorsal pallium. (**A**) Position and cellular heterogeneity of the salamander DP. (**B**) UMAP plot of clusters of glutamatergic neurons from the salamander cortical pallium and glutamatergic amygdala, with highlighted dorsal pallium clusters, data from Woych et al.^52^. (**C**) Heatmap showing differentially expressed genes in superficial- and deep-layer DP neurons, with transcription factors, neuropeptides and genes involved in axon guidance and cell migration/communication highlighted. (**D**) Violin plots showing the expression of *Nts*, *Foxp1*, *Frmd8* and *Nfia* in the superficial- and deep-layer DP neurons. (**E**) Expression of the superficial-layer marker *Nts* and the deep-layer marker *Frmd8* in the salamander telencephalon (E) and dorsal pallium (E’); expression of *Nts* and *Gad2* in dorsal pallium, showing co-expression only in scattered deep layer interneurons (arrowheads) (E”). All markers detected by HCR. (**F**) Expression of transcription factors FOXP1 and NFIA in the salamander dorsal pallium, detected by immunohistochemistry. Scale bar 200 um in overview image (E) and 50 um in magnifications (E’, E”, F). (**G**) Birthdating study design, showing salamander development, and time points of EdU injection. (**H**) Coronal sections at mid-telencephalic level in the juvenile salamander showing the presence of EdU-labeled cells according to the time of injection. White dashed lines depict the pallial-subpallial borders, scale bars 200 um. (**I**) Magnification of the dorsal pallium, showing co-labeling of EdU with *Nts* or *Frmd8* (HCRs). White dashed lines depict the borders of the DP, scale bars 50 um. (**J**) Relative number of EdU+ cells co-expressing *Nts* or *Frmd8* for each injection time. n = 4 for each injection time, except 55a where n = 5, error bars represent standard deviation. Statistical significance was assessed by repeated-measures two-way ANOVA with Geisser-Greenhouse correction, followed by Tukey’s multiple comparisons test (* p < 0.05, ** p ≤ 0.01, *** p ≤ 0.001, **** p ≤ 0.0001) and is indicated above the data point, with the color of the asterisk corresponding to the comparison. Abbreviations: DL, deep-layer neurons; DP, dorsal pallium; dpf, days post fertilization; EGC, ependymoglia cells; INs, interneurons; LA, lateral amygdala; LP, lateral pallium; LV, lateral ventricle; MP, medial pallium; SL, superficial-layer neurons; TEGLU, telencephalic glutamatergic; Tel, telencephalon; VP, ventral pallium

### Pallial layers are generated sequentially during development

We next asked whether the two layers in the anterior DP are generated sequentially over time during development, as are layers in the mammalian cerebral cortex. In *Pleurodeles* larvae, pallial neurogenesis spans approximately three months,^54,55^ and previous work, including intraventricular AAV injections at different developmental timepoints,^49^ suggested that neurons in different positions along the radial axis are generated at different time points during development. We thus designed a birthdating experiment where we administered the thymidine analog EdU to separate cohorts of larvae at different developmental stages, and then assessed EdU labeling in post-metamorphic juveniles (stage 56, within 10 days after metamorphosis), when neurons have integrated into mature circuits^55^ (**Fig. 1G**). Since EdU gets diluted with every cell cycle, only cells that exit the cell cycle around the time of EdU injection will retain EdU labeling at the experimental endpoint. In the pallium, early EdU administration resulted in labeling of superficial neurons, while later EdU injections produced labeling in neurons progressively closer to the ventricle, indicating that salamander pallial neurogenesis follows an outside-in pattern, where early-born neurons are superficial and late-born neurons are deeper. In contrast, neurons in the subpallium, especially the striatum, integrated over the full thickness of the mantle zone, regardless of their birthdate (**Fig. 1H**). Overlaying the expression of the superficial-layer neuron marker *Nts* and deep-layer marker *Frmd8* with the EdU signal revealed that early-born and late-born neurons in the DP are not only distinct in their laminar positioning, but also in their molecular identity (**Fig. 1I**). Quantification suggests that at the population level, neural progenitors in the DP gradually switch from making *Nts*+ superficial-layer to *Frdm8*+ deep-layer neurons around stage 50 (mid-neurogenesis) (**Fig. 1J**). These results reveal a conserved principle of cortical development in amphibians and mammals: the sequential, time-dependent generation of neurons with distinct molecular identities and specific positions along the radial axis.

### Pallial radial glia are multipotent

Two alternative scenarios could explain the sequential generation of salamander superficial- and deep-layer neurons from radial glia progenitors (**Fig. 2A**). One scenario is that DP radial glia are unipotent, with distinct “early” and “late” radial glia types and lineages. “Early” radial glia would be active in early neurogenesis, and cease to proliferate or exist around stage 50, whereas “late” radial glia would be quiescent in early neurogenesis, and activated only around stage 50. Alternatively, DP radial glia may be multipotent, like most radial glia in the mammalian neocortex (part of the mammalian dorsal pallium),^40^ with the same progenitor cell switching from producing superficial- to deep-layer neurons around stage 50.

**Fig. 2.**
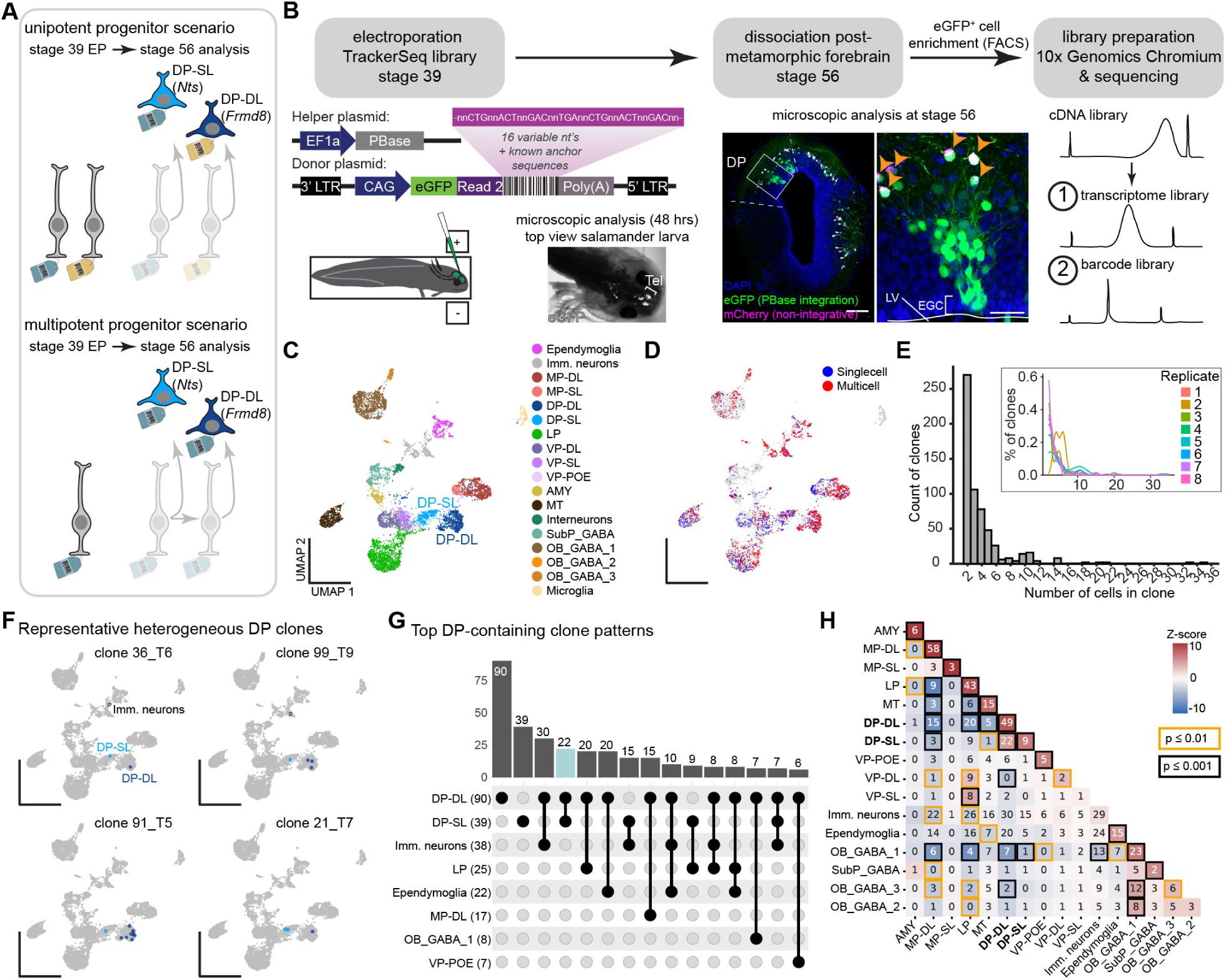
Salamander dorsal pallium radial glia are multipotent. (**A**) Schematic overview of two hypothetical mechanisms to generate neuronal diversity in the DP, testable with barcode-based lineage tracing. Distinct unipotent progenitors could individually give rise to either deep- or superficial-layer DP neurons, leading to different barcodes in either neuron type (top). Alternatively, a single progenitor cell generates multiple neuron types over time during development, leading to shared barcodes in superficial- and deep-layer DP neurons (bottom). (**B**) Experimental setup showing the timeline on top, and experimental details on the bottom. To show that piggyBac integration works in salamanders, the helper plasmid, donor plasmid and a non-integrative pCAG-mCherry plasmid were electroporated at stage 39, and animals analyzed at stage 56. mCherry is expressed only in superficial neurons (early-born), whereas eGFP labels neurons spanning the whole radial axis. Orange arrowheads point to double-positive cells, scale bars are 200 um and 50 um, respectively. (**C,D**) UMAP plot of the TrackerSeq dataset, colored by cell type (C) or TrackerSeq clonal status. Singlecell: barcode detected only in one cell; multicell: barcode detected in multiple cells in the dataset (D). (**E**) Distribution of number of cells in each clone for the complete dataset, or per replicate. (**F**) Examples of clones including both DP-DL and DP-SL neuron types. (**G**) Upset plot showing clonal intersections between groups of DP clusters, with the bar graph on top showing the number of observed intersections (determined inclusively), and the set size per cell type in between brackets on the left. (**H**) Clustermap showing lineage coupling of cells in multicell clones, with the coupling z-score comparing the observed distribution of clonal intersections to many shuffled distributions for each pairwise comparison. Numbers in the clustermap represent the inclusive count of clonal intersections for each pair of cell-groups. Abbreviations: AMY, amygdala; DP-DL, dorsal pallium deep-layer neurons; DP-SL, dorsal pallium superficial-layer neurons; EGC, ependymoglia cells; EP, electroporation; Imm. neurons, immature neurons; LP, lateral pallium; LV, lateral ventricle; MP-DL, medial pallium deep-layer neurons; MP-SL, medial pallium superficial-layer neurons; MT, mitral and tufted cells of the olfactory bulb; OB_GABA_1-3, olfactory bulb GABAergic neurons; SubP_GABA, subpallium GABAergic neurons; VP-DL, ventral pallium deep-layer neurons; VP-POE, ventral pallium post-olfactory eminence; VP-SL, ventral pallium superficial-layer neurons

To test these two hypotheses, we used a barcode-based lineage tracing technique called TrackerSeq, where a barcoded eGFP library is stably integrated in neural progenitors with a PiggyBac transposase, and then barcodes and cellular transcriptomes are retrieved with scRNA-seq to reconstruct lineage relationships.^56^ We electroporated the developing salamander pallium with a TrackerSeq library at stage 39, one of the earliest stages amenable to electroporation of the telencephalon, when the production of superficial neurons peaks (**Fig. 1J, 2B**). Animals were allowed to complete larval development and metamorphosis (3.5 - 6.5 months), after which we confirmed the stable expression of eGFP, and observed groups of spatially-contiguous eGFP+ cells spanning the entire thickness of the pallium (**Fig. 2B**). We then dissociated the electroporated forebrains, FACS-sorted eGFP+ cells, and prepared single-cell sequencing libraries for both the transcriptomes and lineage barcodes (**Fig. 2B**). Eight libraries, each containing traced cells from two animals, were sampled, resulting in the recovery of 8,011 cells. To examine the cellular composition in the dataset, we annotated Seurat clusters using established cell type marker genes^52^ (**Fig. 2C**, **fig. S2A**). To assign clonal relationships, we identified connected components in a graph constructed from TrackerSeq barcode signatures (see **Methods**). This led to the recovery of 1,158 cells, each sharing a lineage barcode with at least one other cell in the dataset, resulting in 285 “multicell” clones (**Fig. 2D,E**, **Tables S1, S2** and **figs. S2-S6**). We counted the number of cells with clonal intersections between annotated groups. Twenty-two clones in the dataset included superficial- and deep-layer neurons (**Fig. 2F,G**). Other clones included postmitotic DP neurons and immature neurons and/or ependymoglia cells (salamander neural stem cells directly derived from radial glia progenitors) (**Fig. 2F-H**). Using these counts, we calculated a z-score for each pairwise set of groups.^57^ This coupling analysis demonstrated a significant clonal relationship between superficial- and deep-layer DP neurons (**Fig. 2H**). Hierarchical clustering of the pairwise correlations between the coupling scores revealed strong correlations between the superficial- and deep-layer DP neurons (**fig. S6A**), further indicating that these cell types originated from the same progenitor during development. These data reveal a second conserved principle of corticogenesis: the generation of superficial and deep layer neurons by multipotent radial glia progenitors.

### Pallial radial glia progenitors are heterogeneous in space and time

Next, we sought to elucidate how superficial- and deep-layer neurons are specified from radial glia progenitors at the molecular level. For this, we built a scRNA-seq atlas of the developing salamander brain (**Fig. 3A, fig. S7**) including eight stages sampled over 2 months of development, from stage 30 (7-day-old embryo) to stage 55a (last stage before metamorphosis), significantly expanding on our previously published dataset.^52^ Our final dataset included 127,488 high-quality cells, grouped in 14 major neuronal and non-neuronal clusters (**Fig. 3B, fig. S7A,B**), which we annotated based on the expression of well-established marker genes (**fig. S7B-D**). Next, we filtered the dataset for telencephalic progenitor cells and neurons, defined as *Foxg1*-expressing cells (**Fig. 3B’,C**), and subsequently retained only cells of the pallium, expressing glutamatergic markers (*Neurog* and *NeuroD* genes) and not GABAergic markers (*e.g. Dlx* genes) (**Fig. 3C’-D’**, see **Methods**). Cells in the UMAP projection were arranged according to the expression of general markers of neuronal differentiation; from cells expressing the radial glia markers *Slc1a3* and *Sox9*, to cells expressing the neural progenitor marker *Sox11*, and then cells expressing mature neuron markers such as the synaptic gene *Snap25* (**Fig. 3G**). To annotate this dataset, we used an iterative supervised learning approach to transfer cell type annotations from our adult cell type atlas^52,53^ to the developmental dataset (**fig. S7E**, see **Methods**), which we validated using established marker genes (**fig. S7F**). The stage 30 dataset was dominated by progenitor cells and immature neurons, and the relative proportion of differentiated neurons increased in subsequent stages. Similar to the mammalian pallium,^58,59^ we observed the first differentiated neurons in the olfactory bulb (OB), followed by ventral regions (amygdala, and ventral pallium) and then more dorsal regions (lateral, dorsal and medial pallium) (**Fig. 3F,G**), consistent with Joven et al.^55^

**Fig. 3.**
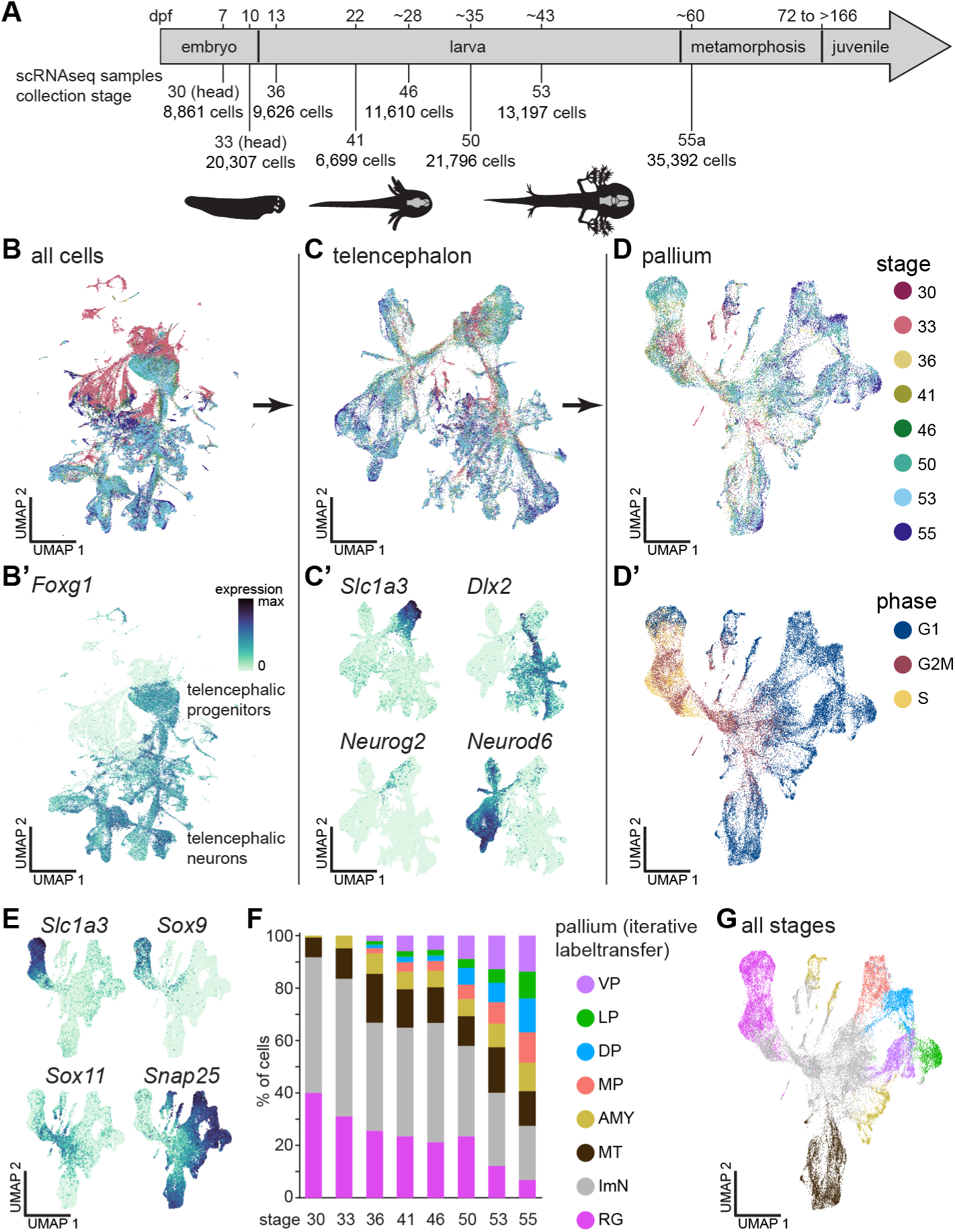
Development of neuronal diversity in the salamander pallium. (**A**) Overview of *P. waltl* embryonic and larval development, with indication of the stages sampled for scRNA-seq. (**B**) UMAP plot of the developmental dataset, colored by sampling stage (B) or the expression of forebrain marker *Foxg1* (B’). (**C**) UMAP plot of the telencephalic subset colored by sampling stage (C) or the expression of *Slc1a3* (progenitors), *Neurog2* (neurogenic pallial progenitors), *Dlx2* (GABAergic lineage) or *Neurod6* (glutamatergic lineage) (C’). (**D**) UMAP plot of the pallial subset colored by sampling stage (D) or cell cycle phase (D’). (**E**) UMAP plots of the pallium, colored by the expression of *Slc1a3* and *Sox9* (progenitors), *Sox11* (neurogenic progenitors) or *Snap25* (neurons). (**F**) Barplot showing the proportion of cells from each sampling stage, colored by labeltransfer annotation. (**G**) UMAP plot of the pallial subset, colored by labeltransfer annotation (marker gene expression shown in fig. S7F). Abbreviations: AMY, amygdala; DP, dorsal pallium; dpf, days post fertilization; LP, lateral pallium; ImN, immature neurons; MP, medial pallium; MT, mitral and tufted cells of the olfactory bulb; RG, radial glia; VP, ventral pallium

Similar to the mammalian cortex, proliferative radial glia in the salamander pallium express *Gfap*, *Sox2*, and *Sox9*,^52,55^ have a soma in the ventricular zone, and a thin fiber extending multiple endfeet to the pial surface (**Fig. 4A,B, fig. S7C**). Studies on the mouse neocortex have identified the temporal molecular states of radial glia progenitors that underlie the generation of early- and late-born neurons.^60^ We thus asked whether salamander radial glia exist in distinct temporal states as well. In our dataset, however, radial glia cells were sampled from different developmental stages, as well as from different spatial domains (pallium and subpallium, and their subdivisions). We reasoned that the transcriptomic diversity of radial glia was the combination of spatial and temporal gene expression signatures, plus other known sources of technical and biological heterogeneity (*e.g.* number of genes and of UMIs per cell, cell cycle phase). Therefore, we first disentangled the spatial heterogeneity of radial glia cells, to then focus on dorsal pallium radial glia and analyze their temporal heterogeneity (**Fig. 4C**). We selected cycling progenitor cells with high expression levels of *Slc1a3*, *Gfap*, *Sox2* and *Fabp7* (also known as BLBP), and absence of *Neurog2* (*i.e.* not yet committed to a neuronal fate) from the pallial dataset (**fig. S8A**). After clustering, we identified and filtered out radial glia cells that exited the cell cycle and acquired an astroglial fate (reduced expression of *Pcna* and *Mcm2* and high levels of astrocyte markers such as *Aqp4*, *Slc13a5*, *Acan*)^55^ (**Fig. 4C, fig. S8B,C**). We then performed a principal component analysis (PCA), after regressing out the effect of known sources of technical and biological variation including the sampling stage (**fig. S8D**), and finally projected the cells on the principal curve that captures the remaining sources of transcriptomic heterogeneity (**Fig. 4C**). A total of 1,783 genes (947 increasing and 836 decreasing) had significant expression variation along this principal curve, with the majority of differentially expressed genes following an expression gradient rather than displaying sharp boundaries. We hypothesized that this principal curve correlated with the medio-lateral position of pallial radial glia, as suggested by the expression of several genes at the two extremes of this axis, including *Wnt8b* and the Wnt antagonist *Sfrp1* (**Fig. 4D**). In salamanders^61,62^ and other vertebrates, Wnt and anti-Wnt signaling molecules define the hem and the anti-hem, two major telencephalic signaling centers at opposite ends of the pallial mediolateral axis. We then clustered pallial radial glia based on these varying genes (**fig. S8E**) to identify major spatial domains. Validation in the stage 50 telencephalon confirmed that these clusters correlated with regional boundaries along the mediolateral axis. *Slit2* and *Dmrt3* were restricted to medial pallium (MP) radial glia, while *Wnt8b* expression extended to part of the DP, although at lower levels. Furthermore, *Meis2* expression was upregulated in DP and peaked in LP, whereas *Sp8* and *Sfrp1* expression spanned the most ventral parts of the pallium (**Fig. 4E, fig. S8F-L**). With these analyses, we were able to identify DP radial glia cells (**Fig. 4C,F**).

**Fig. 4.**
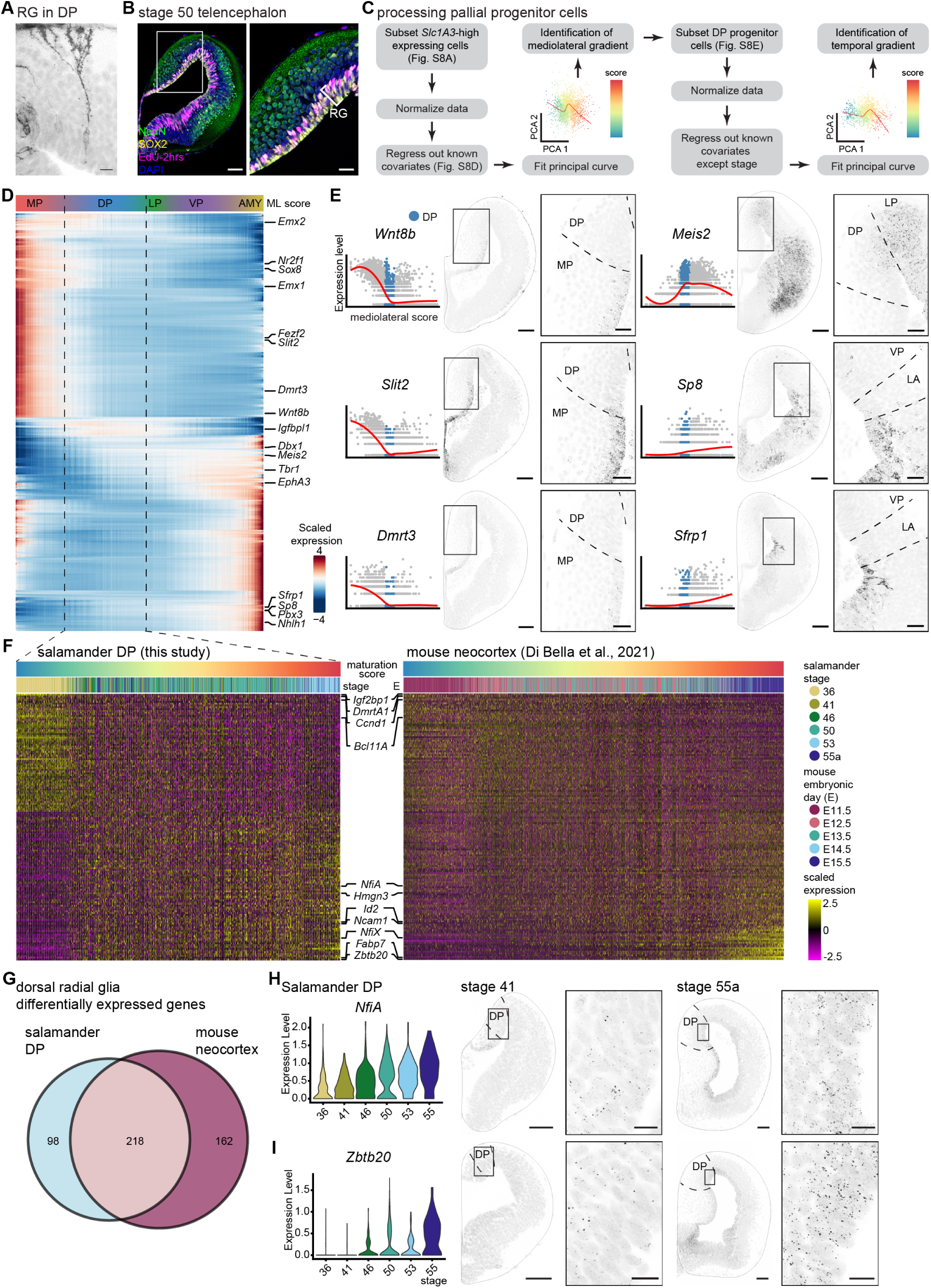
Spatial and temporal patterning of pallial radial glia. (**A**) Visualization of a radial glia cell in the salamander DP through electroporation of membrane-bound GFP at stage 53. (**B**) Coronal section through the telencephalon of a stage 50 larva, injected with EdU 2 hrs before euthanasia, showing immunoreactivity for NEUN (neurons), SOX2 (radial glia) and EdU (proliferative cells). (**C**) Schematic representation of the analysis. (**D**) Heatmap showing differentially expressed genes in pallial radial glia cells, ordered along the principal curve representing the mediolateral score. (**E**) Left: expression (normalised molecules per cell) of canonical markers of mediolateral specification as a function of mediolateral score. The red curve shows local averaging (loess smoothing). Dorsal pallium cells highlighted in blue. Right: gene expression in the telencephalon (HCRs) and magnification. Scale bars in overview 100 um, in magnifications 40 um. (**F**) Heatmaps showing differentially expressed genes in salamander DP radial glia (left) and mouse neocortical radial glia (right, data from Di Bella et al.^63^) arranged along the principal curve representing the maturation score. (**G**) Venn diagram representing differentially expressed genes along the maturation principal curve in salamander and mouse dorsal radial glia (overlapping p-value < 1e-16 after Fisher’s exact test). (**H,I**) Validation of the maturation score, showing expression of *NfiA* and *Zbtb20* in the scRNA-seq dataset (H, violin plots) and in the salamander telencephalon at early (left) and late (right) developmental stages (I). Scale bars in overview 100 um, in magnifications 20 um. Abbreviations: DP, dorsal pallium; EP, electroporation; HCR, hybridization chain reaction in situ hybridization; LA, lateral amygdala; LP, lateral pallium; ML, mediolateral; MP, medial pallium; RG, radial glia; VP, ventral pallium

Next, we analyzed the remaining sources of transcriptomic heterogeneity in DP radial glia. We again performed PCA, after regressing out known sources of variation including the medio-lateral gradient, but excluding sampling stage, and fit a principal curve. The position of cells along this curve correlated with developmental stage, suggesting that this curve captured the maturation of the DP progenitors over time (**fig. S9A**). Hundreds of genes displayed either decreasing or increasing expression along this axis of variation (**fig. S9B**, **Table S3**). We then performed the same analysis on a mouse neocortical dataset^63^ to identify temporal genes in mouse radial glia cells between stages E11.5 and E15.5. Finally, we compared radial glia temporal genes in salamander and mouse. Among salamander-mouse one-to-one orthologs, 316 (188 upregulated and 128 downregulated) genes in salamanders and 380 (168 upregulated and 212 downregulated) genes in mouse showed differential expression across the maturation axis. Of those, a significant subset (218 genes) showed conserved differential expression in mouse and salamanders (**Fig. 4F,G**, **Table S4**). The genes upregulated over developmental time in both species showed enrichment for gene ontology categories such as gliogenesis and glial cell differentiation, while genes downregulated over time in both species were associated with RNA/DNA regulatory functions (**fig. S9C**). Among these shared temporal genes, we found the transcription factors *Nfia*, *Nfix*, and *Zbtb20*, well-known temporal transcription factors in mammalian neural development.^64–69^ Validating these results, HCRs showed higher expression of *NfiA* and *Zbtb20* in dorsal pallium radial glia in late (stage 55a) compared to early (stage 41) salamander larvae (**Fig. 4H,I**). Altogether, these results indicate that in salamander and mouse, dorsal radial glia exist in two conserved temporal states, defined by the expression of a conserved set of temporal genes.

### Intermediate progenitor cells in the developing salamander pallium

The molecular similarity of radial glia progenitors in amphibians and mammals contrasts with the wide transcriptomic divergence of differentiated pallial neurons across species.^52^ To understand how different neuron types can emerge from a conserved developmental blueprint, we sought to compare differentiation trajectories inferred from salamander and mouse datasets. After excluding cells of extra-pallial origin, we applied URD trajectory inference to our entire salamander scRNA- seq pallial dataset (see **Methods**). This approach orders cells according to pseudotime (**fig. S10A**), and then generates a trajectory tree based on transcriptomic similarity.^70^ The resulting tree shows divergence of differentiation trajectories after cell cycle exit, starting from stage 33, and is consistent with current knowledge on pallial regions. Specifically, the first split in the URD tree separates the precursors of OB mitral/tufted cells from the rest of pallial cells, and is followed by a split between dorsomedial and ventrolateral pallial cells, in line with our previous results^52^ and the expression of key regional marker genes (**Fig. 5A-C, fig. S10B**). From an analysis of genes dynamically expressed along the pseudotime axis, we observed that cells first expressed radial glia markers like *Slc1a3*, *Pax6*, *Sox2* and *Sox9*, followed by the Notch receptor *Dll1* and the transcription factors *Nhlh1*, *Insm1*, *Sox4*, and *Eomesodermin* (*Eomes*), which encodes for T-box protein 2 (Tbr2), and finally *Syt1*, *Snap25* and *Slc17a7*, markers of differentiated neurons (**Fig. 5D,E, fig. S10C**). Interestingly, *Dll1, Nhlh1*, *Insm1*, and *Eomes* are key markers for intermediate progenitor cells (IPCs) in the embryonic subventricular zone (SVZ) of the mammalian neocortex.^71,72^ These transient-amplifying progenitor cells are classically considered an amniote innovation and believed to contribute to the expansion of the neocortex.^5,8,73^ However, extending the comparative analysis to the entire transcriptome, we found that the expression of mouse IPC-and radial glia-specific genes was enriched in salamander *Eomes+*/*Slc17a7-* and *Pax6*+/*Sox2*+ cells, respectively (**Fig. 5E,F**). A closer inspection of *Eomes*+/*Slc17a7*- cells in the transcriptomics space revealed that these cells exist in distinct proliferative states (**Fig. 5G**). As described for mouse *Eomes*+ cells,^74^ one subpopulation expressed genes associated with highly proliferative cells (*Top2A*, *Cdk1*, *AurkA*), while a second group appears to be exiting the cell cycle, upregulating differentiation genes such as *NeuroD6* (**Fig. 5G**). HCR revealed *Eomes* mRNA in a subset of cells in the ventricular zone of the salamander pallium, both in early and late developmental stages (**Fig. 5H**). To validate whether these *Eomes* cells are proliferative like mammalian IPCs and as indicated by the scRNAseq data (**Fig. 5C,D**), we detected S-phase cells after EdU administration and stained M-phase cells with a phospho-histone H3 antibody. Both methods revealed the presence of proliferative *Eomes*+ cells in the salamander ventricular zone (**Fig. 5I,J**). Together, these results indicate that *Eomes+* IPCs are not a mammalian innovation, but a conserved progenitor type in vertebrate pallial development.

**Fig. 5.**
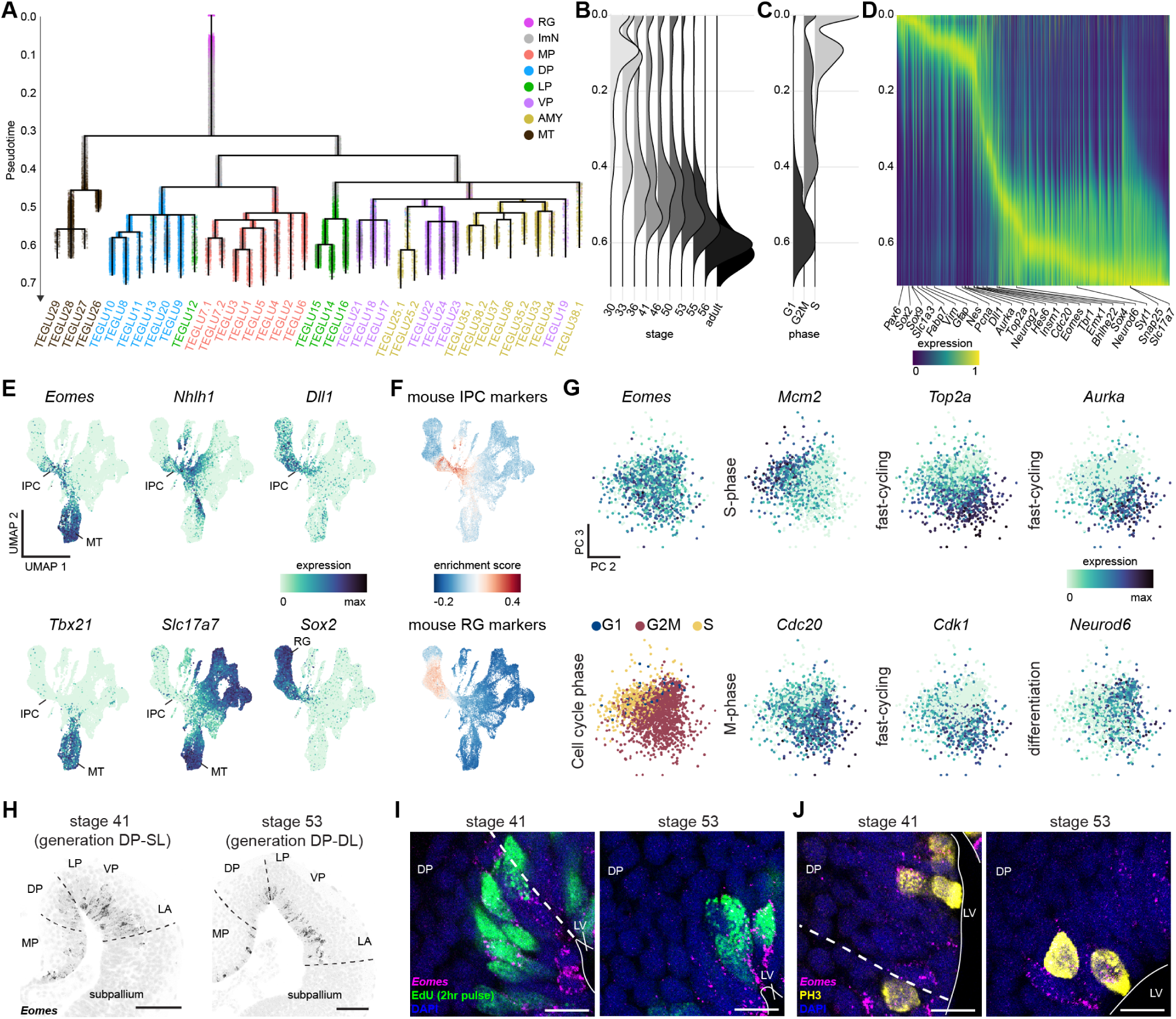
Proliferative *Eomes*+ intermediate progenitor cells in the salamander pallium. (**A**) URD branching tree with 4751 cells ordered by pseudotime, with cells expressing high levels of *Slc1a3* as root, and adult differentiated neurons as tips. Cells are color-coded by their regional identity. (**B,C**) Distribution of cells according to sampling stage (B) or cell cycle phase (C) along the URD pseudotime axis. (**D**) Smoothened heatmap of genes with variable expression in pallium cell trajectories along pseudotime. Only genes expressed in both SL and DL trajectories are shown. (**E**) UMAP plots of the pallium dataset colored by the expression of *Eomes*, *Nhlh1*, *Dll1* (intermediate progenitor cells), *Tbx21* (mitral and tufted cells of the olfactory bulb), *Slc17a7* (glutamatergic neurons), and *Sox2* (RG progenitors). (**F**) UMAP plots of the salamander pallium dataset colored by the enrichment score (Seurat module score) for mouse IPC (top) and RG (bottom) marker genes. (**G**) *Eomes+ Slc17a7-*negative cells (IPCs) in principal components (PC) space, showing subpopulations according to the expression of proliferation and differentiation markers, as well as cell cycle phase. (**H**) Coronal sections through the stage 41 (left) and stage 53 (right) dorsal telencephalon, showing the expression of *Eomes* in scattered cells in the pallial ventricular zone. (**I,J**) Magnifications of the dorsal pallium ventricular zone, showing co-expression of *Eomes* with EdU (2hr pulse, S-phase) (**I**) or PH3 (M-phase) (**J**). Scale bars in H 100 um, in I-J 20 um. Abbreviations: DL, deep layer neurons; DP, dorsal pallium; ImN, immature neurons; IPC, intermediate progenitor cell; LA, lateral amygdala; LP, lateral pallium; LV, lateral ventricle; MP, medial pallium; MT, mitral/tufted cells; PH3, phospho-histone H3; RG, radial glia; SL, superficial layer neurons; St, stage; VP, ventral pallium.

### Conserved projection classes from diverging neuronal differentiation programs

Our comparison of salamander and mouse progenitors revealed striking transcriptomic similarities in radial glia and intermediate progenitors, as well as the conservation of radial glia temporal states. We thus wondered how these progenitors generate diverging sets of glutamatergic neurons in salamanders and mammals. Trajectory analyses show that neurons become increasingly distinct at the transcriptomic level as they differentiate (**Fig. 5A**, ^63,75–77)^, suggesting that major evolutionary divergences may occur when neuronal identity is specified after cell-cycle exit. In line with this, the transcriptomes of radial glia and IPCs are highly correlated in mouse and salamander, but correlations drop once immature and mature neurons are compared (**Fig. 6A**). In salamander, hundreds of genes were differentially expressed along pseudotime in the differentiation trajectories leading to superficial- and deep-layer neurons (**Fig. 6B**). Differentially expressed genes at the split between superficial- and deep-layer trajectories included several transcription factors, such as *Foxp1* (a transcription factor specifying early-born neurons in the mammalian neocortex)^78^ and *Pou3f1* in the superficial-layer trajectory and *Nr3c2*, *Nfix, Nfic*, *Zfpm2*, and *St18* in the deep-layer trajectory (**Fig. 6B-D**). We then compared these trajectory genes with the genes expressed at the first branch-point during mouse neocortical development, which corresponds to the split between intratelencephalic (IT)-projecting and non-IT projecting neurons (**Fig. 6C**).^63,77^ Strikingly, none of the top TFs differentially expressed at the salamander superficial- and deep-layer branch-point showed differential expression at the mouse IT vs non-IT branchpoint (**Fig. 6D**). Moreover, we found that the top TFs distinguishing mouse IT and non-IT trajectories were not differentially expressed in salamander superficial and deep-layer trajectories (**Fig. 6E**). This comparison demonstrates that neurons are specified by different gene regulatory programs in the salamander dorsal pallium and mammalian neocortex, suggesting that evolutionary changes late in pallial development produce divergent neuron types in these two brain regions.^52^

**Fig. 6.**
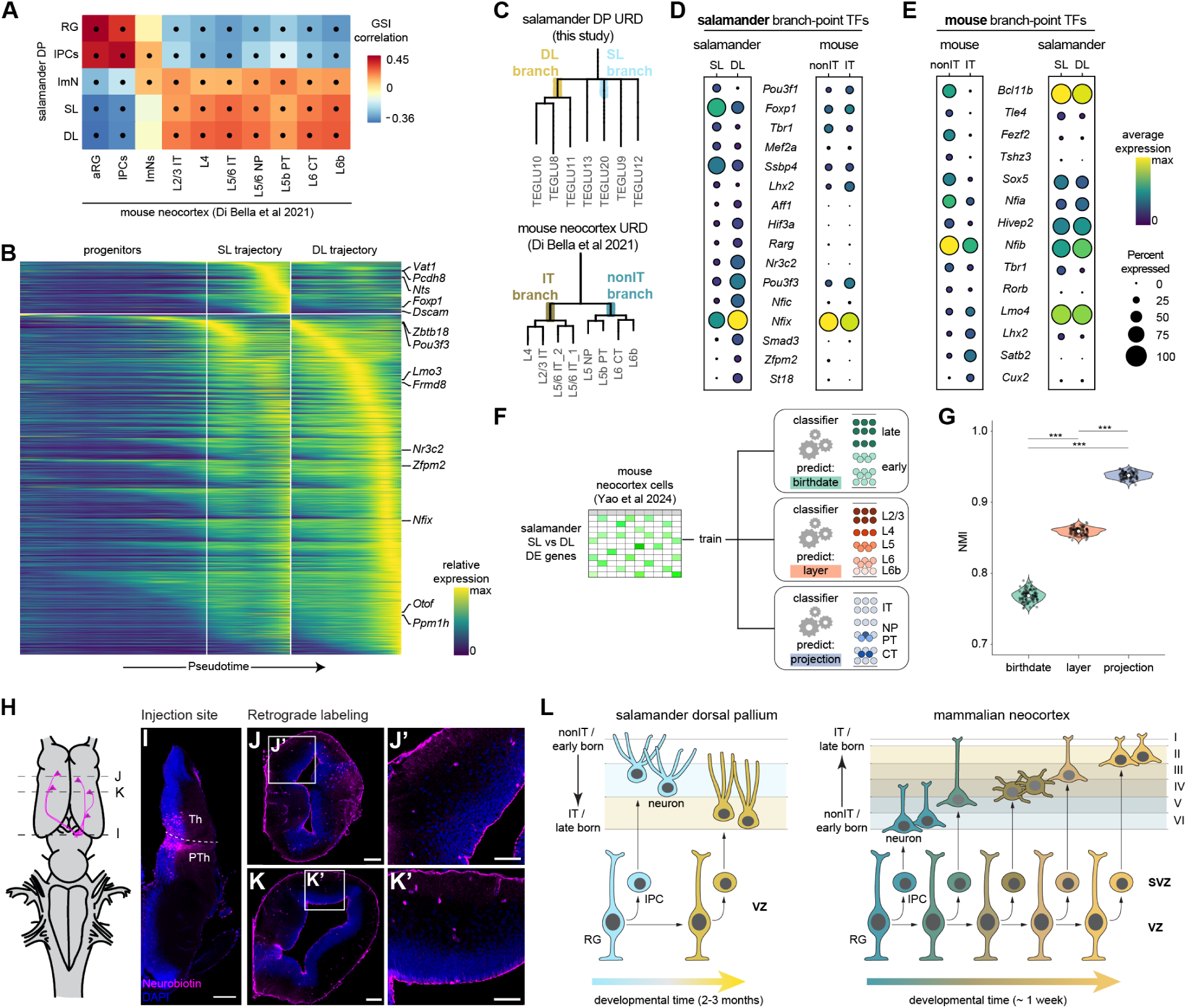
Diverging differentiation programs after cell cycle exit produce conserved classes of pallial projection neurons. (**A**) Gene specificity index correlation matrix of cell types during salamander and mouse neurogenesis showing higher molecular correlations among progenitors, and lower correlations after cell-cycle exit. Dots indicate statistical significance (p<0.05). Mouse data from Di Bella et al.^63^. (**B**) Smoothened heatmap of gene expression in salamander dorsal pallium cell trajectories (**fig. S10A**), showing DE genes between SL and DL trajectories. (**C**) URD tree segments of salamander DP and mouse neocortex (from Di Bella et al.^63^), highlighting branches used for analysis in D,E. (**D**) Left: dotplot of top transcription factors differentially expressed at the URD branchpoint splitting salamander DL and SL neurons indicated in C. Right: expression of the same transcription factors in mouse. (**E**) Left: dotplot of top transcription factors differentially expressed at the URD branchpoint splitting mouse IT and non-IT neurons indicated in C. Right: expression of the same transcription factors in salamander. (**F**) Machine learning classifiers (multinomial logistic regression) were trained to predict alternative classification schemes of mouse neocortical types (birthdate, layer, projection) using only genes differentially expressed across salamander layers. (**G**) Normalized mutual information (NMI) score for each classifier after bootstrapping (NMI=0 indicates no agreement between predictions and ground truth, NMI=1 indicates perfect agreement). p values computed after a Kruskal-Wallis rank sum test and pairwise post-hoc comparisons with Dunn’s test (*** p ≤ 0.001). (**H-K**) Retrograde tracing schematic (H), visualization of the injection site in the thalamus/prethalamus (I) and retrogradely traced cells in the DP (J,K). Magnifications show labeled SL neurons in the DP. Scale bars in overview 200 um, in magnifications 100 um. (**L**) Comparison of neurogenesis in the salamander DP and mouse neocortex. Colors indicate birthdate, cellular morphologies represent divergence of cell identities, and are based on ^24,83^. Abbreviations: aRG, apical radial glia; CT, corticothalamic; DE, differentially expressed; DL, deep layer neurons; DP, dorsal pallium; GSI, gene specificity index; ImN, immature neurons; IPC, intermediate progenitor cell; IT, intratelencephalic; L, layer; NP, near-projecting; NMI, normalized mutual information; PT, pyramidal tract; PTh, prethalamus; RG, radial glia; SL, superficial layer neurons; SVZ, subventricular zone; TEGLU, telencephalic glutamatergic neurons; TFs transcription factors; Th, thalamus; VZ, ventricular zone.

Neuronal identity is defined by the expression of distinct gene modules, each linked to a specific aspect of the neuron’s phenotype or function. Some, but not all gene modules identified in the developing mouse neocortex^77^ were also expressed in salamanders (**fig. S11**). We thus reasoned that salamander and mouse pallial neurons may still share some characteristics because of common features of their early development such as similar birthdates. To explore this possibility, we analyzed mouse neocortical subclasses (as defined by Yao et al.^79^) and tested whether genes differentially expressed between salamander early-born (superficial) and late-born (deep) neurons were sufficient to distinguish between early- and late-born neurons in the mouse. We further asked whether the same gene set could correctly classify mouse neocortical neurons based on their layer or projection identities (**Fig. 6F**). After training machine-learning classifiers (multinomial logistic regression models, see **Methods**) on these data, we assessed their ability to predict neuronal identity across these three axes: birthdate, layer, and projection. Surprisingly, the salamander superficial- and deep-layer specific genes were most effective at predicting projection identity in the mouse, followed by layer and then birthdate (**Fig. 6G**). Consistently, salamander superficial- and deep-layer genes showed enriched expression in mouse neurons with distinct projection identities: salamander superficial-layer markers were enriched in mouse extratelencephalic projection neurons not only in the neocortex, but also in the subiculum, postsubiculum, and retrosplenial cortex, while salamander deep-layer markers were enriched in mouse intratelencephalic-projecting neurons (**fig. S12**).

Tracing and single-cell dye filling experiments in several amphibian species indicate that dorsal pallium neurons are primarily intratelencephalic projection neurons.^80–83^ Prompted by the enrichment of salamander superficial-layer neuron markers in mouse extratelencephalic projection neurons, we performed iontophoretic injections of the bidirectional tracer Neurobiotin in the diencephalon of adult salamanders, to verify the existence of pallial neurons projecting to extratelencephalic regions. Retrogradely-labeled cell bodies were detected in several telencephalic regions, with cell bodies in the dorsal and medial pallium mostly restricted to the superficial layer (**Fig. 6H-K**). Given that superficial-layer neurons are a small fraction of all DP neurons, it is likely they were missed in previous studies, which document only intratelencephalic projections from DP.^52,80–83^ Together, these data indicate that salamander superficial- and deep-layer neurons largely correspond to extra- and intratelencephalic projection neurons, respectively. In summary, our results show that neuronal birthdate, laminar position, molecular identity, and projection type are tightly linked in the salamander dorsal pallium, just as they are in the mammalian cerebral cortex, revealing a shared developmental logic underlying circuit assembly in both regions (**Fig. 6L**).

## Discussion

In this work, we characterized the development of the amphibian dorsal pallium at single-cell resolution. Our findings overturn the longstanding assumption that pallial multipotent radial glia, temporal patterning, IPCs, and layers are innovations in amniotes or mammals. We identify conserved principles of corticogenesis, and uncover what truly distinguishes cortical development in mammals from other vertebrate species (**Fig. 6L**).

A first conserved principle of cortical development is the generation of distinct glutamatergic neuron types in a stereotyped temporal sequence by multipotent radial glia progenitors. In the neocortex, these progenitors produce clones that span the entire radial axis.^84^ Hippocampal radial glia progenitors are also believed to be multipotent and generate spatially-clustered clones.^41^ Our TrackerSeq analysis indicates that salamander radial glia cells are multipotent and generate both superficial and deep layer neurons. Like in mice,^60^ salamander radial glia exist in distinct molecular states over time and, importantly, a conserved set of genes distinguishes early and late radial glia cells in both species. This finding suggests that early and late radial glia states are homologous and that temporally-patterned pallial neurogenesis traces back to tetrapod ancestors. An emerging question is whether the diversity of radial glia temporal states correlates with the total number of neuron types and layers generated. In insects, where neuroblasts exist in discrete transcriptional states over time, the diversification of these temporal states may be necessary for the evolution of new neuron types.^85^ In contrast, our data suggest that the evolution of a complex cell type repertoire in the mammalian neocortex was unlikely to have occurred through changes of temporal patterning, because salamander and mouse temporal states are largely comparable. Finally, we find that these conserved radial glia states do not track absolute time, but relative time. In salamanders, pallial neurogenesis spans 2-3 months, while in mice it lasts only about a week. This points to the existence of global mechanisms that align and scale the *tempo* of neurogenesis to the development of the rest of the organism.

A second conserved principle of cortical development is the amplification of the neuronal output of radial glia by intermediate progenitor cells. While the evolutionary conservation of radial glia cells is well established, the evolution of IPCs remains heavily debated.^3,5,7,9,50,51^ In the mammalian neocortex, IPCs delaminate from the ventricular surface and aggregate in a distinct band of cells called the subventricular zone (SVZ), where they divide and produce neurons.^5,59^ With the identification of an SVZ in the pallium of turtles and birds, it has been proposed that IPCs are an amniote innovation supporting the generation of large pallia, particularly in birds and mammals.^7,9,86^ Challenging this view, a Tbr2+ (*Eomes*) expressing SVZ was discovered in the shark pallium.^51,87^ Scattered non-ventricular mitoses have also been reported in amphibians, but *Eomes*/Tbr2 expression in these putative IPCs could not be confirmed.^48,88^ Our transcriptomic analysis solves this conundrum by demonstrating that salamander *Eomes*+ progenitors express the battery of genes that defines mammalian IPCs, in line with recent results in chicken^89^ and sharks^50^. These cells are mitotic and committed to a neuronal fate, suggesting the presence of indirect neurogenesis. Together, these data indicate that *Eomes*-expressing IPCs existed in the pallium of jawed vertebrate ancestors, most likely to increase the neuronal output of radial glia cells. The flexible deployment of this progenitor type, culminating with their delamination into an SVZ multiple times independently across vertebrates, may have enabled the expansion of selected pallial regions without fundamental modifications of radial glia progenitors.

A third and central principle of cortical development is the distribution of neuron types according to a birthdate gradient along the radial axis, producing a tight association of neuronal birthdate with molecular identity, laminar position, and projection identity.^44^ This is the case not only in the neocortex, but also in the hippocampus and piriform cortex, where neurons with different birthdates settle into distinct sublayers and exhibit distinct transcriptomes, connectivity, and functions.^27,31,32,34^ Temporally-patterned neurogenesis, present in other parts of the nervous system and across species, is not the exclusive developmental strategy for generating layered structures (*e.g.* the superior colliculus and cerebellum)^90,91^. The finding that birthdate is linked to neuronal identity and layering in the salamander pallium as it is in mammals suggests that this mechanism for pallial layer generation was already present in tetrapod ancestors. This hypothesis is further supported by data from other vertebrates. In turtles, lizards, frogs, and zebrafish, pallial neurons born at different developmental times are arranged into a gradient along the radial axis.^29,42,43,47,48^ In the gecko^15^, alligator^30^, and turtle^33^ dorsal cortex, superficial and deep neurons have distinct gene expression profiles. Notably, in the turtle dorsal cortex, superficial neurons are early-born and extratelencephalic-projecting, whereas deep-layer neurons are late-born and intratelencephalic-projecting,^29^ resembling the salamander dorsal pallium. These similarities extend to the transcriptomes of salamander and turtle superficial and deep-layer neurons,^52^ suggesting that these two layers are homologous across species and were present in tetrapod ancestors. To date, no evidence in fish directly indicates a link between neuronal birthdate, transcriptomic identities, and projections. However, zebrafish early-born pallial neurons congregate into a nucleus (Dc) that has a distinct transcriptomic identity;^92^ these neurons are extratelencephalic projecting, whereas late-born neurons are mostly intratelencephalic types.^47^ These observations suggest that the dichotomy between early-born extratelencephalic-projecting and late-born intratelencephalic-projecting neurons identities may even extend to vertebrate ancestors.

A striking difference between mammals and other vertebrates is then the direction of the neurogenesis gradient along the radial axis: outside-in (early-born neurons in superficial layers, late-born neurons in deep) in turtles^29,42^, lizards^15,43^, amphibians^48,49^, and fish^47^ as opposed to inside-out (early-born neurons in deep layers, late-born neurons in superficial) in mammals (**Fig. 6L**). Pallial lamination may have evolved convergently multiple times in vertebrates, leading to dissimilar radial distributions of early- and late-born neurons. Alternatively, if early- and late-born neurons are homologous across tetrapods, the mammalian inside-out pattern of corticogenesis could have arisen through the inversion of an ancestral outside-in pattern. In this scenario, a critical mammalian innovation was the emergence of a new mode of radial migration. Changes in Wnt signaling in immature neurons may have enabled later-born neurons to gain the ability to migrate past earlier-born neurons, as indicated by experiments in reptiles.^93^ Cajal-Retzius cells, which in mammals regulate radial migration through Reelin signaling, are present in non-mammalian species, but their increased abundance and higher *Reelin* expression in the mammalian pallium may have also contributed to the emergence of radial migration.^94^ The inversion of the corticogenesis gradient, together with changes of thalamocortical connectivity,^6,95^ may have relieved constraints for the further diversification of late-born intratelencephalic projection neurons, supporting the expansion and specialization of cortico-cortical connections in mammalian ancestors.

Unlike in reptiles and amphibians, the mammalian cerebral cortex includes areas with multiple layers and greater neuronal diversity. Our findings indicate what kinds of developmental innovations may have led to the generation of more neuron types in mammals. We show that, while early steps of corticogenesis rely on largely conserved progenitor types, the differentiation cascades that unfold after cell cycle exit differ markedly between the mouse neocortex and salamander dorsal pallium. We propose that the evolutionary divergence of neuronal differentiation programs was instrumental for the increase of neuronal diversity and layers in mammals - although we cannot fully rule out that these differentiation programs stem from molecular heterogeneities in radial glia or IPCs. This divergence is so profound that prior transcriptomic comparisons of adult pallial neurons could not identify clear one-to-one homologies across species.^33,52,61^ Our developmental perspective sheds new light on this staggering neuronal diversity. We propose that neuronal birthdates identify two ancestral classes of pyramidal neurons in the dorsal pallium - early-born extra-telencephalic and late-born intra-telencephalic projection neurons - and that diversification within each of these classes produced the plethora of neuron types that populate not only the mammalian neocortex, but also adjacent areas such as the entorhinal cortex and the subicular complex.^35,52^ Comparisons across vertebrates are needed to understand how this neuronal diversification contributed to functional innovations in cortical circuits.

## Acknowledgments

We thank the Columbia University ICM team for the excellent animal care, all members from the Tosches lab for critical input throughout the project, and Darcy Kelley for feedback on the manuscript. We thank Chao Feng and Elena Dvoretskova (Mayer lab) for the generation of the TrackerSeq libraries and fruitful troubleshooting, Saeed Tavazoie for access to FACS sorting, Lina Habba for help with HCRs, and Susan Brenner-Morton for the FOXP1 antibody. Computing resources were provided by Columbia University’s Shared Research Computing Facility (NIH grant 1G20RR030893-01 and NYSTAR contract C090171).

## Funding

This work was supported by the National Institute of Health (grants R35GM146973 to MAT and 1RM1HG011014 to RS), the Rita Allen Foundation (MAT), the Chan Zuckerberg Initiative (MAT), the McKnight Foundation (MAT), a DFG project grant 549328218 (CM), an EMBO Long-Term Fellowship ALTF 874-2021 (AD), the National Science Foundation (Graduate Research Fellowships to EG and LKVPL), the Anusandhan National Research Foundation (ANRF/ECRG/2024/006839/LS to SC), and the TiH Foundation (TIH0018-002 to SC).

## Author contributions

AD and MAT designed the study. AD, EG, and JW collected scRNA-seq data. AD, SC, CL, EG, MAT analyzed scRNA-seq data with input from RS and CM. AD and JW performed histology and imaging. AD and AOG performed birthdating analysis. AD performed injections and electroporations. AD and LKVPL optimized and implemented FACS. PA produced and analyzed axonal tracing data. AD and MAT wrote the manuscript, with edits from CL and SC and feedback from all other authors. CM and RS provided critical feedback on the project. MAT supervised the whole study.

## Competing interests

In the past 3 years, RS has received compensation from Bristol Myers Squibb, ImmunAI, Resolve Biosciences, Nanostring, 10x Genomics, Parse Biosciences and Neptune Bio. RS is a co-founder and equity holder of Neptune Bio.

## Data and materials availability

The scRNA-seq data produced for and used in this study have been deposited in the Gene Expression Omnibus (GEO) with accession numbers GSE308588 (stages 30-53), GSE309006 (stage 55a) and GSE309390 (TrackerSeq). Code used for the analysis of TrackerSeq data is available on https://github.com/mayer-lab/TrackerSeq.

## Materials and Methods

### Animals/sampling

*Pleurodeles waltl* embryos, larva and adults were obtained from breeding colonies established at Columbia University. The animals were maintained in an aquatics facility at 20-25 ℃ under a 12L:12D cycle.^96^ All experiments were conducted in accordance with the NIH guidelines and with the approval of the Columbia University Institutional Animal Care and Use Committee (IACUC protocols AC-AABF2564 and AC-AABL1550). Embryos and larvae were staged according to Gallien et al.^97^.

### EdU administration

Larvae were deeply anesthetized in 0.02% MS-222 and placed on their back in a petri dish or Sylgard mold. A single EdU injection (2.5 or 5 mg/mL stock solution in sterile 0.9 % saline with FastGreen at a 1:30 ratio) was administered intraperitoneally (50 mg/kg of total body weight), using a glass capillary needle and pressure injection at developmental stage 37, 41, 46, 50, 53 or 55a. Larvae were allowed to complete development and metamorphosis, and were sacrificed at stage 56, specifically 0-2 weeks after completion of metamorphosis. To minimize animal-to-animal variability, larvae were maintained in controlled environmental conditions throughout the experiment.

### DNA constructs

Some expression vectors used in electroporation experiments were purchased from Addgene: pCAG-mCherry (gift from Jordan Green, Addgene plasmid # 108685) and pCAG-mGFP (gift from Connie Cepko, Addgene plasmid # 14757).

### TrackerSeq library preparation and validation

The TrackerSeq plasmid library was produced following the library production protocol from Bandler et al.^56^. In brief, a 16-base oligonucleotide randomer was cloned into a linearized pCAG-EGFP vector using 12 reactions of HiFi assembly. Subsequently, the reconstituted plasmids were desalted and concentrated. Then, 8 reactions of 4.5 uL barcoded plasmid were mixed with 50 uL NEB-beta 10 electrocompetent cells and electroporated in a 5 mm cuvette at 1.95 kV, 200 W, 25 uF with 3.8-4.0 ms. Transformed bacteria were recovered for 15 minutes and then plated onto 15 cm LB + Amp + Sucrose plates for recovery of individual colonies, then scraped and cultured in 5L LB + Amp + Sucrose media for 3 hours (until OD = 4.8), followed by maxiprep and concentration.

### Electroporation

Glass capillary needles were pulled and then broken open using fine forceps (needle tip diameter ranging 5-15 um), backfilled with mineral oil, and then connected to a Nanoject III injection system (Drummond), and filled with plasmid solution. Right before injection, larvae were deeply anesthetized in 0.02% MS-222 and stabilized in a Sylgard mold. The needle was inserted through the skin, in the right lateral telencephalic ventricle, and plasmid solution was pressure injected. At stages 39 or 53, 120 nL or 250 nL plasmid solution was pressure injected at 10 nL/s or 15 nL/s, respectively. For pCAG-mGFP electroporation, a plasmid DNA solution of 1.5 ug/uL was prepared. For TrackerSeq experiments, a DNA plasmid solution of TrackerSeq library (final concentration of 0.8 ug/uL) and pEF1a-pBase (PiggyBac-transposase; a gift from R. Platt) (final concentration of 0.7 ug/uL) was prepared. Fastgreen was added to the plasmid preps at a 1:10 ratio for visualization of the injection site.

Immediately after injection, larvae were placed sideways in a freshly prepared 2% agarose carved mold, filled with ice cold PBS, with two additional carved molds for the electrodes at a distance of 7 mm to one another. The electrode molds were positioned either parallel, or in a 40 degree angle relative to the mold for the larva. Electroporations were performed towards the dorsal walls of the ventricle, with a BTX electroporator (ECM830), using round 5 mm diameter platinum plate electrodes attached to tweezers (Nepagene, CUY650P5), similar to what was described in Joven et al.^55^. 5 50 ms unidirectional pulses with a 1 s interval were applied at 38 V or 46 V at stages 39 or 53, respectively, after which the larvae were allowed to recover.

For TrackerSeq experiments, larvae were screened 48-120 hours after electroporation, and those with a bright GFP positive telencephalon were grown to complete larval development and metamorphosis (3.5 - 6.5 months after electroporation).

### Neurobiotin retrograde tracing

Iontophoretic injections were conducted by first anesthetizing salamanders in 0.1% MS222, followed by transcardiac perfusion with oxygenated Amphibian Ringer’s solution (96 mM NaCl; 20 mM NaHCO_3_; 2 mM KCl; 10 mM HEPES; 11 mM glucose; 2 mM CaCl_2_; 0.5 mM MgCl_2_), as previously described Woych et al.^52^. The bidirectional tracer Neurobiotin (VectorLabs SP-1120) was diluted to 5% w/v in oxygenated Amphibian Ringer’s solution immediately before each injection and mixed with FastGreen dye (5% w/v, Sigma-Aldrich F7252) to confirm successful tracer injection. This tracer solution was manually loaded into a microneedle (diameter ∼10 μm) pulled from capillary glass (1.5mm, Sutter P87). The needle was then mounted to a microelectrode holder (WPI MEH35W15) with a silver wire making contact with the tracer solution. The positive terminal was connected to the needle, which was mounted on a manual micromanipulator for injection (Narishige MM3), and the negative terminal was placed under the head of the salamander at the level of the diencephalon, contralateral to the injection site (in order to draw the tracer into the tissue area of interest). Using anatomical landmarks, the needle tip was inserted into the thalamus, and a positive current was applied with parameters 3 μA, alternating 7 sec on, 7 sec off, for 20 min (according to ^98^) using a Midgard Precision Current Source (Stoelting, 51595). Once the injection was completed, the salamander heads were transferred to ice-cold oxygenated Amphibian Ringer’s solution to incubate for 24-30 hours. The brains were then extracted and fixed overnight in 4% PFA at 4°C and processed as described below. Labeling was scored manually and localization of Neurobiotin signal was determined using molecular and anatomical landmarks.

### Tissue processing for (whole mount) IHC and HCR

Animals were first deeply anesthetized by immersion in 0.02% MS222 for larvae, or 0.1% MS222 for post-metamorphic juveniles or adults. From developmental stage 50 onwards, animals were transcardially perfused with ice-cold DEPC phosphate-buffered saline (DEPC-PBS) to remove blood from the brain. Only post-metamorphic animals were additionally perfused with ice-cold 4% paraformaldehyde (PFA) in DEPC-PBS. Brains were dissected, and fixed overnight in 4% PFA in DEPC-PBS at 4 °C. Brains were then dehydrated (40%, 60%, 80%, 100%, 100% methanol in PBS-DEPC, 15 min each at room temperature (RT)) and incubated in 100% DCM overnight at RT, washed twice in 100% methanol and stored at -20°C until further processing.

HCRs with and without EdU detection were performed as described in Jaeger et al.^49^. Specifically, brains were stained as whole-mount preparations, embedded in 4% agarose in Tris-HCl (500 mM, pH7.0) and sectioned at 70 um using a vibratome. Sections were then incubated in DAPI in Tris-HCl for 30 min, and mounted in Fluoromount-G® Mounting Medium (SouthernBiotech) or DAKO fluorescent mounting medium (Agilent Technologies). HCR-3.0- style probe pairs against *Dmrt3*, *Eomes*, *Frmd8*, *Gad2*, *Meis2*, *Nfia*, *Nts*, *Sfrp1*, *Slit2*, *Sp8*, *Wnt8b*, *Zbtb20* were designed using insitu_probe_generator^99^ and ordered from IDT, and 6 pmol was added to the probe hybridization buffer (Molecular Instruments) for incubation. For co-staining with antibodies SATB1 (1:50, abcam ab51502), PH3 (1:500, Sigma 06-570) or GFP (1:500, abcam ab13970), the primary antibody was added to the amplification buffer with hairpins, and the secondary antibody being either goat anti-mouse IgG Alexa 488 antibody, donkey anti-chicken IgY Alexa 488 (1:500, Jackson ImmunoResearch), or goat anti-rabbit IgG Alexa 546 (1:500, Invitrogen) to the overnight wash in 500 mM Tris-HCl, similar to what has been described for the GFP antibody in Jaeger et al.^49^. For the detection of Neurobiotin, Streptavidin conjugated to Alexa Fluor 647 (1:1000, Invitrogen S32354) was added to the amplification buffer.

Immunostainings were performed on 70 um vibratome sections. Briefly, sections were permeabilized in PBS supplemented with 0.2% Triton X-100 (PBST) for 30 min at RT, then blocked in blocking buffer (2.5% BSA, 2.5% sheep serum, 50 mM glycine in PBST) for 30 min at RT and incubated with primary antibody solution (10 mM glycine, 0.1% H2O2 in PBST) for 1-3 nights at 4 °C. Tissue sections were then washed 3 x 15 min in PBST at RT, incubated with secondary antibody solution (DAPI 1:1000 in PBST) for 2 hrs at RT, and again washed for 3 x 15 min before mounting in Fluoromount-G® Mounting Medium (SouthernBiotech) or DAKO fluorescent mounting medium (Agilent). Primary antibodies used are rabbit anti-FOXP1 (1:40000, custom antibody generated by Susan Brenner Morton ^100^), mouse anti-NFIA (1:10, DSHB PCRP-NFIA-2C6), mouse anti-NEUN (1:500, Sigma-Aldrich MAB377), rabbit anti-SOX2 (1:500, abcam ab97959), and secondary antibodies goat anti-rabbit IgG Alexa 546 or goat anti-mouse IgG Alexa 647 (1:500, Invitrogen). The NFIA antibody required antigen retrieval, which was performed after permeabilization by incubation in a 10 mM sodium citrate pH6 buffer in a 70 °C water bath for 30 min, followed by 2 5 min washes in PBST at RT.

### Confocal microscopy and EdU quantification

All images were acquired using a Zeiss LSM800 confocal microscope, and processed in Fiji. HCR images presented in grey were generated by setting the channel colors to Grays and inverting the LUTs of the HCR and DAPI channels separately, and then overlaying both channels in Adobe Photoshop and adjusting the transparency of the DAPI channel. For quantification of the molecular identity of EdU cells, sections across the anterior-posterior axis of the anterior dorsal pallium (until the ventricle of the two telencephalic hemispheres merges in the coronal section, 8-14 hemispheres per animal) were imaged as tiled z-stacks (6 images at 4 um interval). Using the Cell Counter Plugin in Fiji, all EdU cells in the dorsal pallium were manually counted and scored for expression of *Nts* and *Frmd8*. Cells were considered positive for EdU when the intensity of the label was above background levels, regardless of the pattern within the nucleus. Cell proportions were then calculated, and data presented as mean ± SD (n = 4 animals analyzed for each stage, except for stage 55a for which n=5).

### Head or brain dissociation and single-cell capture

To increase the comprehensiveness of our previously published single-cell RNA-sequencing developmental dataset^52^, we dissociated 40 heads from stage 30, and 25 heads from stage 33 embryos. In addition, we dissociated the telencephalic vesicles of an additional 6 stage 50, and of 6 stage 53 larvae, and the telencephalon and diencephalon of 15 stage 55a larvae. The latter received an intracerebroventricular AAV-PHP.eB virus injection for driving the expression of eGFP under the CAG promoter at either stage 37, stage 41 or stage 51 as described in Jaeger et al.^49^. Dissociation was performed according to Woych et al.^52^, with a small modification for the stage 30 and 33 heads, which were only incubated for 15 min in the enzymatic dissociation cocktail. All dissociated cells were multiplexed using 4 distinct Cell Multiplexing Oligos (10x Genomics 3’ CellPlex Multiplexing kit) per stage, prior to GEM formation.

### Sample collection for TrackerSeq libraries

We separately collected the telencephalon, and the diencephalon and midbrain of electroporated brains from salamander juveniles (right after metamorphosis) and pooled tissue chunks from two animals before performing papain dissociation according to Woych et al.^52^ with minor modifications to recover as many cells as possible. All filters were pre-wetted and post-washed with 1-5 mL fresh carbogenated calcium-free Hibernate A media (BrainBits), centrifugation was performed at 500 x g, and after the density gradient centrifugation step, the cell pellet was resuspended in calcium and magnesium free PBS supplemented with 1% BSA and transferred to FACS tubes. All tubes for filtering the cells, and the FACS tubes were coated overnight with 2% BSA in calcium and magnesium free PBS at 4 °C. To enrich electroporated cells, we performed fluorescence activated cell sorting using a Bio-Rad S3e Cell Sorter (ProSort software version 1.6.) with a 100 um nozzle. Non-eGFP-expressing brain tissue from control siblings was used as a negative control for excluding background fluorescence and to set the gate on forward scatter. The positive diencephalon and midbrain samples were used to set the lasers and gates of the eGFP population. Sorted cells (between 2300 and 13000 cells per sample) were collected in bulk in coated low-profile PCR 8-strips with temperature control set to 4 °C. Cells were processed on the 10x Genomics Chromium platform.

### Analysis single-cell RNA sequencing data for the developmental dataset

#### Read alignment

For all newly acquired samples, reads were aligned to the combined *Pleurodeles waltl* transcriptome^52^ using Alevin (Salmon v1.6.0)^101^. The transcriptome was supplemented with itr-eGFP sequences for aligning the stage 55a libraries. Library type was set to automatic, and keepCBFraction was set to 1, whereas all other parameters were set as default. Developmental libraries were multiplexed using the 10x Genomics 3’ CellPlex Multiplexing kit, and multiplex tag reads were assigned using Alevin as well.

#### Data QC and filtering

Count matrices obtained after alignment with Alevin were imported in the R package Seurat (version 4.3.0.1)^102^ to generate individual Seurat objects. The objects were first demultiplexed independently based on the identity of the Feature Barcode oligonucleotides introduced using the 10x Genomics 3’ CellPlex Kit, and only singlets were retained. These objects were then filtered to remove droplets with low UMI counts (less than 700 reads per cell for all libraries except stage 30, which was filtered to remove cells with UMI counts below 2000), and high mitochondrial content (more than 15% mitochondrial genes per cell). Datasets were then pooled in steps depending on when the samples were generated and further filtered using Seurat 4.1.0. Specifically, after Louvain clustering, low-quality cells were removed using a Support Vector Machine (SVM) classifier (R package e1071) as described in Tosches et al.^33^. Low-quality clusters (number of genes per cell below average of the dataset, percentage of mitochondrial genes above average of dataset, and absence of cluster-specific marker genes) were first identified, and 10% of cells from these clusters, together with 10% of cells from the rest of the dataset were used to train the classifier. The trained SVM classifier then identified low-quality cells across each dataset, which we removed. All SVM-cleaned, pooled datasets were finally merged, also including our already published dataset^52^ to obtain a final Seurat object of 127,488 cells.

#### Clustering

The full 127,488-cell dataset was initially clustered and analyzed using the R package Seurat 4.1.0^102^. CellCycleScoring was applied, after which the raw counts were normalized using SCTransform,^103,104^ while regressing the data for the number of UMIs per cell, developmental stage, percent mitochondrial genes, G2M and S scores. The top 3000 variable genes were used and a principal component analysis (PCA) was performed, after which the first 180 principal components were used for clustering and UMAP embedding. The default clustering method was used, with a resolution of 1.5. Clusters were then annotated based on the expression of marker genes, and label transfer (see below). This dataset was analyzed in more detail and at higher resolution for specific clusters. To study the competence of dorsal progenitor cells, pallial progenitors were subsetted in steps based on marker gene expression (presence of *Slc1a3*, *Foxg1*, *Emx1*, absence of *Dlx1/2*, *Gsx2*, *Nkx2.1*, *Ascl1*, *Otx2*, *Cdh6*). Clusters with highly expressing *Slc1a3*, *Gfap*, *Sox2* and *Sox9* cells were then used for spatial and temporal analysis of radial glia progenitors (see below), while clusters with highly expressing *Eomes* cells were analyzed for the presence of intermediate progenitors. For the latter, raw counts were normalized using SCTransform, regressing for the number of UMIs per cell, percent mitochondrial genes and developmental stage (not cell cycle score). 45 principal components were used for clustering and UMAP embedding at a resolution of 0.5. To study neuronal differentiation of the pallium, we implemented a two-step process. First, telencephalic progenitors were selected from the progenitor object, and telencephalic neurons from the full dataset based on a combination of marker genes (*Foxg1*, *Neurod2*, *Neurod6*, *Dlx5*, *Dlx6*, *Snap25*) and labeltransfer (see below). Raw counts from the telencephalic cells were normalized using SCTransform, regressing for number of UMIs per cell, developmental stage, percent mitochondrial genes, G2M and S scores, 150 principal components were used for clustering and UMAP embedding at a resolution of 1.5. From the telencephalic dataset, a dataset including only pallial cells was extracted using a combination of marker genes and labeltransfer (**Fig. 3F,G, fig. S7F**). Raw counts were again normalized using SCTransform, regressing for number of UMIs per cell, developmental stage, percent mitochondrial genes, G2M and S scores, 125 principal components were used for clustering and UMAP embedding at a resolution of 1.5.

#### Label transfer

To annotate the developmental clusters, Seurat’s Label transfer was performed using the adult *Pleurodeles* dataset.^52,53^ Cluster TEGLU7 in medial pallium was split in two after subsetting and running FindAllMarkers, since part of this cluster (TEGLU7.2) expresses *Nts* at high levels with cells located in the superficial MP, while neurons in cluster TELGLU7.1 do not express *Nts* (**fig. S1C,D**). The Label Transfer algorithm identifies transfer anchors between two data sets, allowing comparison of cell type identities between the two. To increase the matching of cell types, Label transfer was performed iteratively. Hereto, the full adult dataset^52,53^ was used as first reference, and the stage 55a dataset as first query. Subsequently, the annotated stage 55a dataset was used as a reference for identifying cell identity in the stage 53 object, and so on.

For all transfers, the function FindTransferAnchors was run using dims = 20, and k.anchor = 5, and the function TransferData was run using k.weight = 20.

#### Heterogeneity of pallial progenitor cells

We first subsetted the pallial dataset to clusters with high *Slc1a3* expression (**fig. S8A,B**). Within this subset, we identified a cluster of cells with a strong astroglial signature (high expression of *Acan*, *Aqp4*, and low expression of *Mcm2* and *Mcm5* (**fig. S8A-C**)). This cluster represents radial glia cells that are differentiating into ependymoglia, and therefore was removed for the rest of the analysis. To characterize the heterogeneity of pallial progenitor cells, we calculated the variance of the normalized data. The data was processed using SCTransform vst.flavor=”v2”.^103^ We calculated the first 50 principal components using the RunPCA() function in Seurat^102^ and then retained the 14 first principal components which captured at least 1% variation each. To identify which factors are associated with each principal component, we calculated pearson correlation between the principal components and various technical and biological factors associated with each cell including cell cycle score “G2M.Score”, “S.Score”, total RNA content (“nCount_RNA”), number of genes expressed in each cell (“nFeature_RNA”), percentage of mitochondrial and ribosomal genes expression (“percent.mt”, “percent.ribo”) of the cell, and developmental stage of the sample (“Stage”) (**fig. S8D**).

To adjust for differences arising from technical and biological factors we re-ran SCTransform with vst.flavor=”v2” and regressed these factors by using vars.to.regress = c(“G2M.Score”, “S.Score”, “nCount_RNA”, “nFeature_RNA”, “percent.mt”, “percent.ribo”, “Stage”) and also re-ran PCA. Next, we fit a principal curve which represents an ordered ‘spatial trajectory’ for each cell based on their expression profile using the principal_curve function in the princurve R Package ^105,106^ using options smoother = ‘lowess’, f = ⅓ and stretch=2). We defined a ‘mediolateral score’ as the length of the arc from the beginning of the curve to the point where the cell projects onto the curve. Since there is no directionality associated with the curve, we defined an explicit starting point based on the expression of *Wnt8b*, such that its expression is negatively correlated with the ’mediolateral score’. *Wnt8b* is a canonical marker for medial pallium, confirmed in the developing *Axolotl* brain using spatial transcriptomics^61,62^ and as such, its expression should be high in the medial cells. To identify genes that vary across the mediolateral axis, we used the fitGAM function in the tradeSeq package^107^ using nknots = 3. To limit the analysis to highly informative genes, we ran fitGAM with mediolateral scores as the pseudotime with only genes that had a residual variance of at least 0.5 after running SCTransform. To assign explicit spatial labels to the progenitors, we performed a hierarchical clustering using the genes that are differentially expressed across the mediolateral trajectory. We used the hclust() function with method=”ward.D2” to perform hierarchical clustering in R; the tree was cut at h=350 to obtain 10 clusters and isolate the dorsal pallium progenitors.

#### Temporal patterning of dorsal pallium progenitor cells

To characterize the genes involved in temporal patterning of dorsal pallium in salamanders, we used a scoring strategy similar to our medio-lateral score. Following our spatial analysis and hierarchical clustering, we subsetted out the dorsal pallium progenitors (**fig. S8E**). To reprocess the subsetted DP population with SCTransform, we regressed out non-temporal factors that contribute to heterogeneity using vars.to.regress = c(“G2M.Score”, “S.Score”, “nCount_RNA”, “nFeature_RNA”, “percent.mt”, “percent.ribo”). We fit a principal curve as previously described (**fig. S9A**). We defined a ‘temporal score’ as the length of the arc from the beginning of the curve to the point where the cell projects onto the curve. For processing the mouse neocortical progenitors, we used SCTransform with vars.to.regress =c(“G2M.Score”, “S.Score”, “nCount_RNA”, “nFeature_RNA”, “percent.mt”, “percent.ribo”). To identify genes expressed differentially across the temporal trajectory, we used the fitGAM function with temporal scores as the pseudotime. Temporally variable genes were identified with a log fold change of 2 and adjusted p-value < 0.05. To identify conserved temporal genes across salamander and mouse, we subsetted the list of temporally variable genes to only include one-to-one orthologs that are expressed in both the species and have a residual variance of at least 0.5. Gene ontology analysis was performed for upregulated and downregulated genes separately using the clusterProfiler package^108^ with the universe genes as the list of one-to-one orthologs that had a residual variance of at least 0.5 in both mouse and salamanders.

#### Analysis of developmental trajectories

To reconstruct transcriptional trajectories that underlie specification and differentiation of pallial neurons, we used the R package URD (v1.1.1)^70^. The pallial object generated previously contains some excitatory neuronal cell types present in the telencephalon, that originate from extra-pallial territories (e.g. *Sim1*+/*Otp*+ neurons in the amygdala, excitatory neurons in the septum, and some mitral and tufted cells in the olfactory bulb). Since their complete trajectory cannot be reconstructed, we removed these neurons for the URD dataset. To increase the number of mature cells, we added differentiated pallial neurons from our adult object. In total, we analyzed 2996 radial glia cells, 13483 immature neurons, and 13840 mature neurons spanning the AMY, VP, DP, LP, MT, and MP clusters.

First, we calculated a diffusion map using the calcDM function in URD with knn=200 and using sigma.use=NULL which automatically determines the most optimal sigma. We assigned a subset of cells with high expression of *Slc1a3* as the root cells after removing the gliogenic cells expressing high *Aqp4*. We then used the root cells and simulated diffusion from root cells to all other cells to calculate the pseudotime. To simulate diffusion, we used the floodPseudotime function in URD with n=50 simulations and using minimum.cells.flooded=2 so that the simulation stopped if 1 or no cells were newly visited in a given iteration. The tips were defined to be the cells that were previously annotated to be mature cells (assigned a TEGLU* label) using our label transfer approach. We then used the pseudotimeWeightTransitionMatrix function with optimal.cells.forward = 40, max.cells.back = 80, pseudotime.direction = “<” to identify the slope and inflection point of the logistic function which is used to bias the transition probabilities using pseudotimeWeightTransitionMatrix function. To simulate random walks on the cell-cell graph from the root cells to the tip, we used the simulateRandomWalk function performing 10,000 random walks per tip (n.per.tip = 10000). To generate the dendrogram layout, we used the buildTree function with divergence.method = “preference”, cells.per.pseudotime.bin = 25 and bins.per.pseudotime.window = 8. To create the forced directed layout, we used the treeForceDirectedLayout function with num.nn = 100, cut.unconnected.segments = 2.

Genes differentially expressed between URD segments were computed using the FindAllMarkers function in Seurat, and filtered to retain genes with adjusted p-value <0.001 that were expressed in at least 20% of at least one segment, and with abs(pct$1 - pct$2)>0.15. From this list, we then identified genes with specific expression in SL or DL trajectories by computing genes expressed in segment 6 (SL) but not in segments 45, 44, 5, 9, and 37 (DL), and vice versa (see fig. S10A for segment labels). The heatmap in Fig. 5D shows all genes with differential expression along pseudotime except these SL and DL-specific trajectory genes, whereas Fig. 6B shows only the SL and DL-specific trajectory genes.

To generate the heatmaps in Fig. 5D and 6B, gene expression was smoothened by computing moving averages along pseudotime, followed by spline fitting (function geneSmoothFit in the URD package). Genes were then ordered according to the relative peak of expression along pseudotime, following the approach described in Raj et al.^109^. For each gene, smoothened expression data were scaled to the max expression for that gene before plotting.

To identify genes with differential expression at URD branchpoints, we used the FindMarkers function in Seurat. For mouse, we used the neocortex developmental data and URD analysis from Di Bella et al.^63^, comparing segments 15 (IT branch) and 17 (non-IT branch). For salamander, we took segment 45 (the segment in common between the three anterior DL cell types) and cells in segment 6 (SL cells) with the corresponding pseudotime values. We filtered these lists of differentially expressed genes to retain only transcription factors (mouse and salamander one-to-one orthologs) with avg_logFC >1 (mouse) or avg_logFC >1.2 (salamander), and expressed in at least 25% of cells in one of the two segments.

To analyze the expression of gene modules in salamander developmental trajectories, we used the gene modules computed by Gao et al.^77^ from a developmental cell type atlas of the mouse visual cortex. For each gene module, salamander-mouse one-to-one orthologs were identified using eggnog mapping. After this step, only modules with at least 5 genes were retained. Expression enrichment for each module was computed using the AUCell_run function in the AUCell package^110^.

#### Gene specificity index correlations of salamander and mouse cells

Correlations between developing salamander and mouse cells were computed following an approach described in Tosches et al.^33^. Briefly, average gene expression from log-normalized data from the developing mouse neocortex (data from ^63^) and the developing salamander dorsal pallium were computed using the AverageExpression function in Seurat. Next, for each species, we computed differentially expressed genes (FindAllMarkers function in Seurat, logfc.threshold=1.5). From these lists of differentially expressed genes, we identified the salamander-mouse one-to-one orthologs, and then took the union of the two lists (5743 genes). Gene specificity matrices for salamander and mouse data were computed as described in Tosches et al.^33^, and then pairwise Spearman rank correlations between cell classes were computed from these matrices. Statistical significance was assessed with a permutation test, after shuffling gene expression values 1000 times across cell classes.

#### Classification of mouse neocortical cells using salamander SL and DL marker genes

To assess how gene signatures derived from SL (TEGLU20) and DL neurons (TEGLU8, TEGLU10, TEGLU11) in salamander DP discriminate cellular identities in the mouse cortex, we first identified differentially expressed genes between salamander SL and DL neurons using the FindAllMarkers function from the Seurat R package (parameters: test.use = “roc”), and then filtered the list to keep only genes expressed in at least 30% of cells of at least one of the two groups, with abs(avg.logFC) > 1.1 and with a ROC score below 0.4 or above 0.6. This resulted in 258 marker genes, of which 148 expressed in SL and 110 in DL. Next, we identified mouse one-to-one orthologs of these genes using eggnog-based mapping. This resulted in 131 SL genes and 88 DL genes. We then used these to compute gene module enrichment in mouse cells. From the Allen Brain Cell atlas^79^, we selected all telencephalic glutamatergic subclasses (defined by the expression of *Foxg1* and *Slc17a6* or *Slc17a7*) and downsampled the dataset to 200 cells per cell subclass. Next, we used AUCell^110^ to compute expression enrichment of SL and DL markers for each subclass in the mouse (**fig. S12**).

Next, we filtered the mouse Allen Brain Cell atlas dataset to keep only neocortical glutamatergic cell “subclasses” (according to the Allen Brain Cell atlas taxonomy) and the expression values of the mouse one-to-one orthologs of salamander SL and DL marker genes (computed as described above). Count data were log-normalized and then scaled. Next, we trained multinomial logistic regression models with L2 regularization (alpha = 0) using the glmnet package’s cv.glmnet function (family = “multinomial”, 20-fold cross-validation). Models were trained to predict one of three classification schemes in mouse metadata: “layer”, “birthdate”, and “projection”. These labels were assigned according to the metadata of the Allen Brain Cell atlas (for birthdate, L5 and L6 neurons were “early born”, and the rest “late born”; for projection, we used IT, ET, CT, and NP labels in the original dataset). We evaluated model performance using a bootstrap approach with 100 iterations. In each iteration, a subset of cells from the mouse were resampled with replacement using sample(), and the remaining out-of-bag (OOB) cells were used for testing. Normalized Mutual Information (NMI; computed via NMI from aricode) was calculated for each replicate. To assess whether classification performance differed significantly for the three sets of labels, we used the Kruskal–Wallis rank sum test, a non-parametric alternative to one-way ANOVA that does not assume normality, followed by pairwise post hoc comparisons using Dunn’s test (FSA::dunnTest()) with Benjamini–Hochberg correction for multiple testing.

### TrackerSeq analysis

#### TrackerSeq transcriptome library processing

The transcriptomic libraries were aligned as described for the developmental libraries above. Count matrices obtained after alignment with Alevin were imported in the R package Seurat (version 4.3.0.1^102^) to generate individual Seurat objects. The objects were first filtered to remove droplets with low UMI counts (less than 700 reads per cell) and low nFeatures (less than 2000 genes per cell). Doublets were identified using DoubletFinder^111^, where the expected number of doublets was estimated assuming a 0.8% doublet formation rate per 1000 cells recovered, adjusting for homotypic doublets based on cluster composition. Parameter optimization was performed by sweeping across principal components 1-30 to select the optimal pK value per library based on bimodality coefficient metrics. Additionally, cells expressing both glutamatergic (*Slc17a6/7*) and GABAergic (*Gad1/2*) genes were annotated as doublets. Cells classified as doublets were subsequently excluded from downstream analyses. Datasets were then pooled, and cells with high mitochondrial content (>10%) were removed.

#### TrackerSeq clustering

The filtered dataset was clustered and analyzed using the R package Seurat (version 4.3.0.1^102^). CellCycleScoring was applied, after which the raw counts were normalized using SCTransform^103,104^, while regressing the data for the number of UMIs per cell, percent mitochondrial genes, G2M and S scores. The top 3000 variable genes were used and a principal component analysis (PCA) was performed, after which the first 35 principal components were used for clustering and UMAP embedding. The default clustering method was used, with a resolution of 1.5. Non-neuronal and GABAergic neuron clusters were then annotated based on the expression of marker genes,^52^ while Seurat clusters of glutamatergic neurons were first subsetted. This subset was clustered with the same parameters as above, except for the resolution which was increased to 3, and clusters were annotated based on the expression of marker genes.^52^ Cells with a poor transcriptomic signature were removed at this point (after checking that these cells are not drivers of clonal relationships).

#### Estimation of TrackerSeq library diversity

From the concentrated TrackerSeq plasmid library, PCR amplification of the variable barcode sequences was performed using a P7-containing Read2 and a P5-containing custom reverse primer, at a low cycle count (∼18) and purified using 2-sided selection for the ∼330 bp length product with SPRI magnetic beads. Over 6.7M NGS reads were performed on the library, with almost 5.4M unique barcodes detected. The barcodes were read-extracted and clustered using Bartender v1.1 bartender_extractor_com and bartender_single_com^112^ to remove putative barcode reads that were likely the result of PCR amplification or NGS sequencing errors using a hamming distance of 2. This resulted in at least 2.6M unique barcode sequences, indicating that the TrackerSeq library used has minimum diversity well into the millions.

#### Preprocessing of TrackerSeq Barcodes

Dedicated TrackerSeq barcode libraries were generated from an aliquot of the 10x Genomics cDNA by amplification with indexed P5 and P7 primers as in Bandler et al.^56^. The resulting NGS fastq files were paired, quality-filtered, and assigned to unique cell barcodes using the UMI-tools ‘extract’ and ‘whitelist’ funmum hamming distances. Complete barcodes with valid UMIs and cellbcs, and including the fixed sequence handle and spacers were clustered using umi-enabled Bartender v1.1 bartender_single_com^112^ to remove putative barcode reads that were likely the result of PCR amplification or NGS sequencing errors. This was done using a hamming distance of 5, since the fixed handle sequence increases minimum hamming distances. Resulting raw barcode sequences and their cluster centroids (the consensus sequence from clustering) were mapped as a dictionary for cluster scores with a quality score of <0.8 (indicating that for each UMI of an erroneous barcode, ∼3x the number of centroid UMIs are observed). Unmapped barcodes were retained in the dataset as unique signatures, which would cause only splitting of clones, and not result in bundling errors, or false positives. Unique combinations of lineage barcodes, cellbcs, and UMIs were aggregated, and UMIs above this threshold counted for each unique pair of lineage barcode and cellbc. The barcode data was filtered for minimum number of reads = 5 and minimum number of UMIs = 3.

To determine clonal membership of cells, we have employed a network-based analysis which treats cells as nodes containing barcodes, with edges connecting each cell that share at least a single barcode. Each cell-cell edge receives a weight according to the following equation:

(*Intersection*(LBC_scell1_, LBC_scell2_)/(LBCs-_cell1_ + LBCS_cell2_)) * 2

The metric scores the shared profile of barcodes present in both cells similar to a Jaccard distance, but with increased penalty for cell pairs where one member of the pair has few barcodes and the other has many barcodes. For each collected dataset, these scores are graphed as a histogram (e.g., counts of cell-cell edges with a given weight) (**fig. S3**).

After edge weighting, all connected components (i.e., cells that are connected by edges) within the network are determined for stepwise weight thresholds from 0 to 1, where the edges with weights below the threshold are removed from the network, filtering membership of connected cell nodes. With this method, cells connected by the strongest lineage barcode signatures are preserved, while those with partial signatures such as those caused by PCR amplification errors or repeat sampling of barcodes are omitted. Since the noise level of each experimental dataset depends on the accumulation of technical noise over multiple experimental steps (**fig. S3**), it is necessary to select a cutoff threshold for each dataset.

Heuristic indicators such as clone number and size statistics were used to mitigate the effect of noise present in unfiltered barcode signatures to optimize recovery of true positives while excluding false positives. Over-filtering the networks would lead to a loss of multi-cell clones that provide meaningful biological signals, as shown by the counts of multicell clones present in the data. Therefore, for each dataset in the TrackerSeq experiment, we selected the minimum threshold value where the clone size distribution was stable. The selected thresholds are shown in each of supplementary **figures S3, S4 and S5**. Connected networks from each dataset were assigned as clones. In addition, as a strict control experiment, we filtered the networks for only perfect lineage signatures (i.e., similarity score = 1) to ensure that commonly observed lineage relationships are retained in the data. We confirmed that this strictly filtered dataset supports the claims made in this paper despite a fractional loss of lineage clones.

#### Analysis of clonal information in transcriptomic data

Clone assignments from the selected threshold levels were mapped into the Seurat transcriptome data. 1988 cells out of 2556 detected in the lineage matrix were successfully mapped. We counted the number of clonal intersections and the number of cells from those intersections (**Supplementary Tables 1 and 2)**.

To evaluate the significance of cell lineages shared across annotated groups, we calculated a coupling z-score for each pairwise comparison of cell categories according to previous work^56,57^ with 5000 permutations of data shuffling on the pooled lineage data from 8 replicates. Z-scores shown in Figure 2F describe the number of standard deviations between the observed clonal counts and the aggregate mean of shuffled permutations. Positive scores indicate observation of more shared clones than chance, indicating positive lineage coupling, while negative scores indicate fewer observations than chance, indicating negative lineage coupling. Z-scores > 2.58 have equivalent p-values ≤ 0.01 and > 3.29 with p ≤ 0.001.

Correlations using np.corrcoef on the between-group scores were performed along with hierarchical clustering using ‘average’ from scipy.cluster (**fig. S6A**).

Some lineage tracing experimental replicates had substantially higher clone counts and coverage of cell types within traced cells (**fig. S2D**). We performed z-scoring using the same method on data for each replicate, then converted these z-scores into percentiles so that inter-sample differences were normalized. For cell-cell pairs that had coverage across all experimental replicates, we used pingouin.rm_anova to perform repeated-measures ANOVA testing between the pairwise lineage counts data for dataset vs. cell-pair assignment. The significant result, f = 3.21 equating to p = 0.015, demonstrated that the variance between cell-cell pair groups was significantly greater than the variance across replicates for each pairwise comparison (**Fig. 2H**).

## Notes

### Summary of Updates

The revised version includes a major restructuring of the Introduction and changes to the Discussion

