## Supplementary figures for "A conserved logic for the development of cortical layering in tetrapods"

*Supplementary figures S1-S12*

*Supplementary tables S1-S4*

### *Supplementary figures*

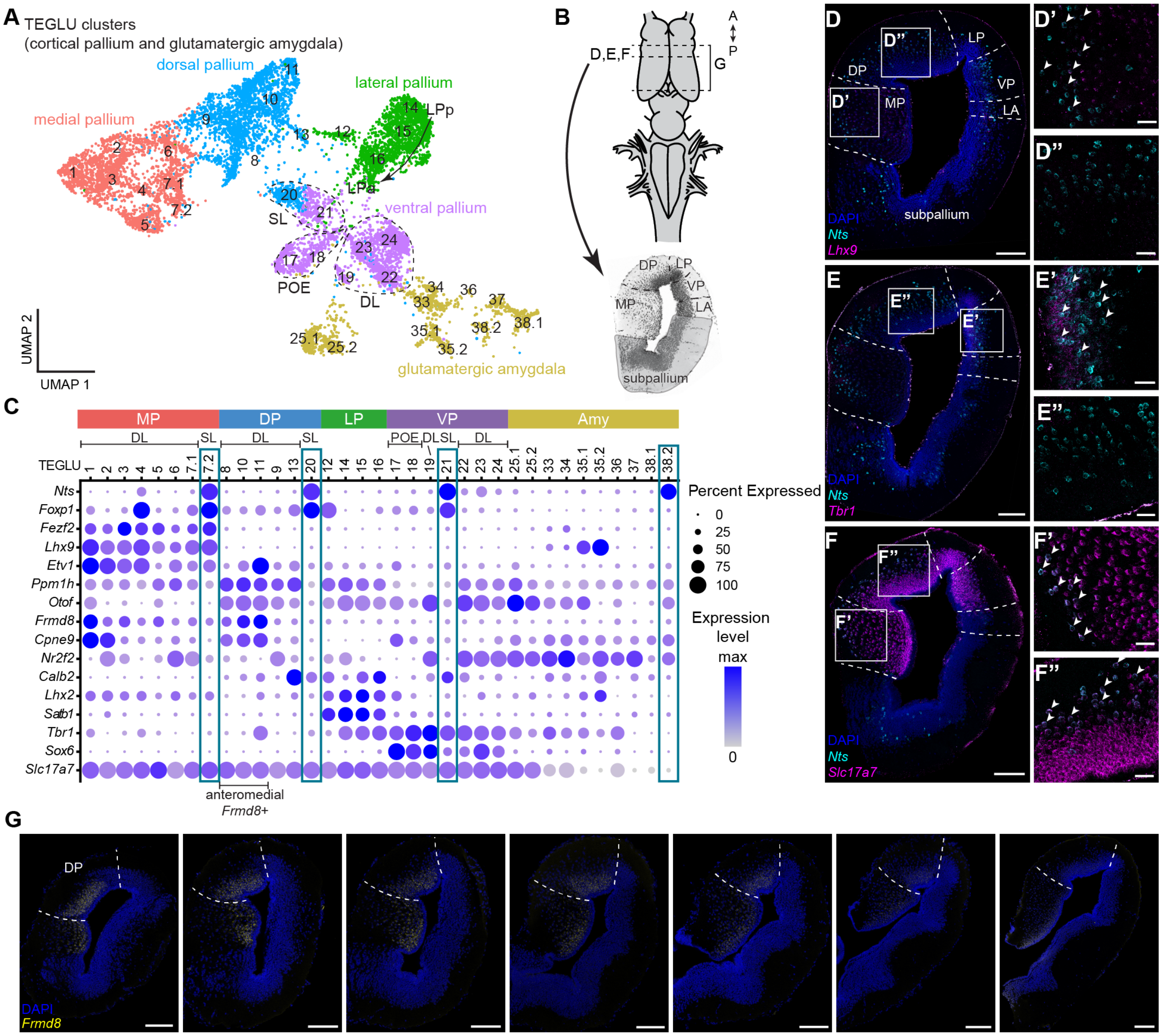


**fig. S1. Glutamatergic *Nts*- or *Frmd8*- expressing neurons in the pallium.**

(**A**) UMAP plot of clusters of glutamatergic neurons from the salamander cortical pallium and glutamatergic amygdala, colored by their spatial location. (**B**) Top: dorsal view on the salamander brain with dashed lines indicating coronal sectioning planes for D-G. Bottom: coronal section through the salamander forebrain at the level of D,E,F showing the pallial domains. (**C**) Dotplot showing TEGLU clusters in the cortical pallium and pallial amygdala. *Nts*-expressing neurons are highlighted, with marker genes for each pallial domain suggesting their localization. (**D-F**) Expression of *Nts* and either *Lhx9* (MP), *Tbr1* (VP) or *Slc17a7* (cortical pallium) in the salamander telencephalon. Dashed lines indicate boundaries between pallial domains, arrowheads point towards a selection of double-positive cells. Scale bar 200 um in overview image and 50 um in magnifications. (**G**) Expression of DP-DL marker *Frmd8* across the anterior-posterior axis, showing a decreased expression in posterior DP. Scale bar 200 um.

Abbreviations: A, anterior; Amy, amygdala; DL, deep-layer; DP, dorsal pallium LP, lateral pallium; MP, medial pallium; P, posterior; POE, post-olfactory eminence; SL, superficial-layer; TEGLU, telencephalic glutamatergic; VP, ventral pallium

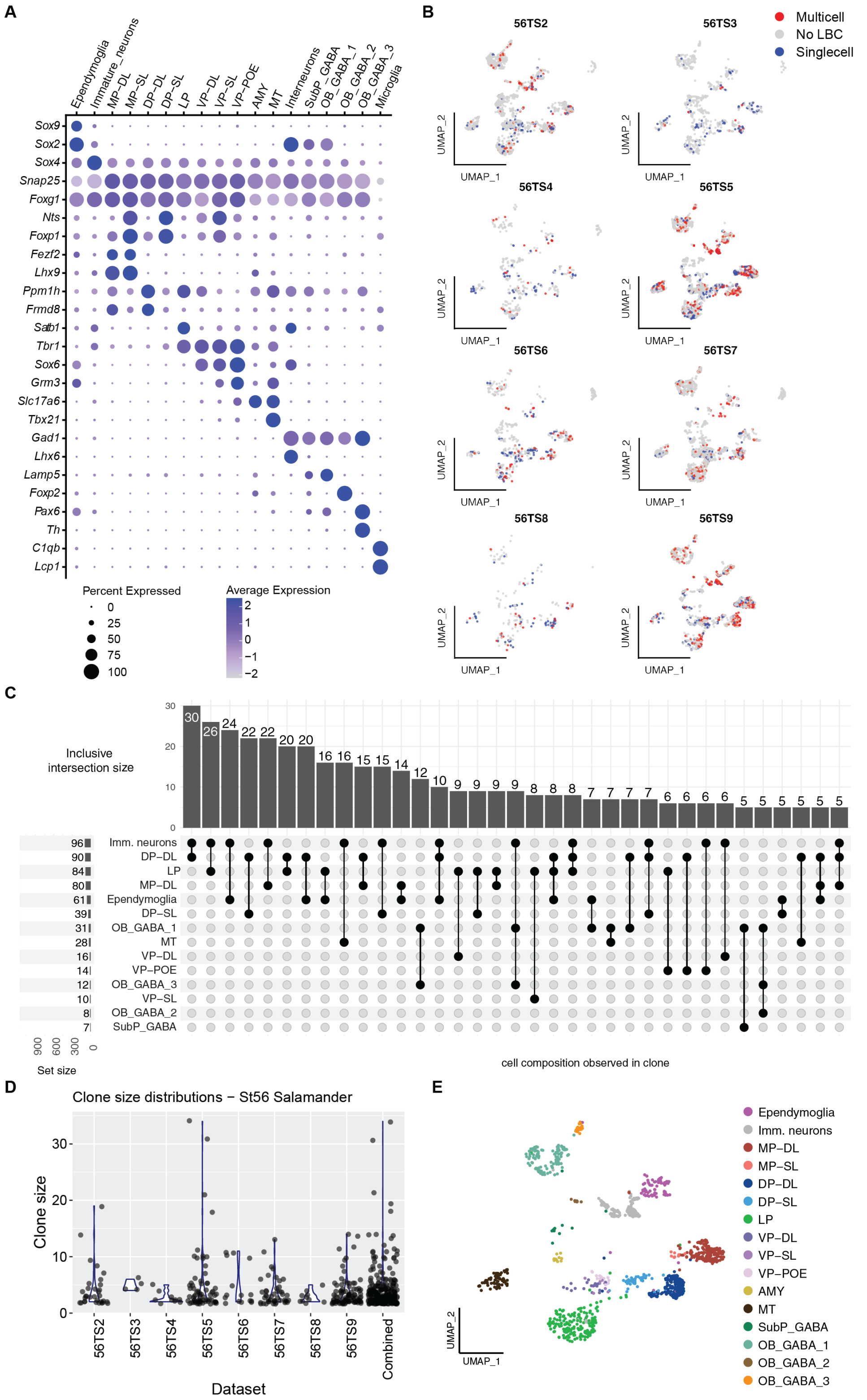


**fig. S2. TrackerSeq dataset characterization**

(**A**) Dotplot showing key marker genes expressed in the clusters annotated in Fig. 2C. (**B**) Single-cell UMAPs showing cells from each experimental replicate, colored by TrackerSeq clone type. Grey = no TrackerSeq lineage barcode (LBC) detected, blue = single-cell clone, red = cell is a member of a multi-cell clone. (**C**) Upset plot of clonal intersections by annotated cell type category, observed at least 5 times and counted inclusively. For each lineage pattern shown in the grid, the bar on top indicates the number of individual clonal intersections where this pattern was observed. Clones may be counted more than once if they match multiple patterns shown. (**D**) Distribution of TrackerSeq multicell clone sizes shown by experimental replicate. (**E**) Single-cell UMAP showing only cells belonging to a multicell clone, colored by cell type.

Abbreviations: AMY, amygdala; DP-DL, dorsal pallium deep-layer neurons; DP-SL, dorsal pallium superficial-layer neurons; Imm. neurons, immature neurons; LP, lateral pallium; MP-DL, medial pallium deep-layer neurons; MP-SL, medial pallium superficial-layer neurons; MT, mitral and tufted cells of the olfactory bulb; OB_GABA_1-3, olfactory bulb GABAergic neurons; SubP_GABA, subpallium GABAergic neurons; VP-DL, ventral pallium deep-layer neurons; VP-POE, ventral pallium post-olfactory eminence; VP-SL, ventral pallium superficial-layer neurons

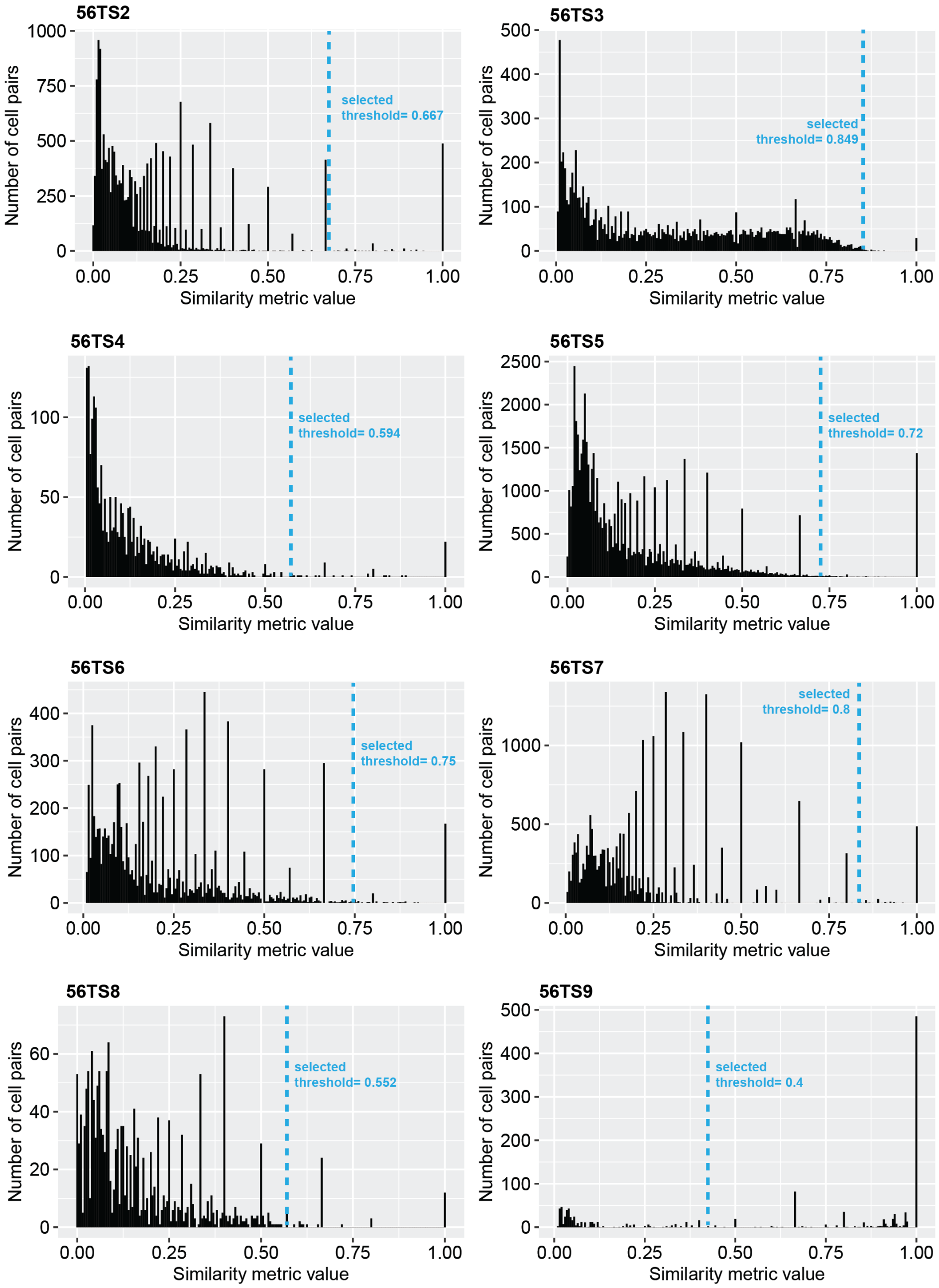


**fig. S3. TrackerSeq barcode histograms**

TrackerSeq cell-lineage barcode network edge weight histograms for each TrackerSeq experimental replicate generated in the study. Each pair of cells is connected by an edge in the network, which is weighted based on the average of common vs. total barcodes observed in each cell. This weight is shown along the x-axis, with the total number of edges for each weight shown on the y-axis. The blue dashed line indicates the edge weight that was used to filter the network. Only edges with weights above the selected threshold were retained for clonal assignments.

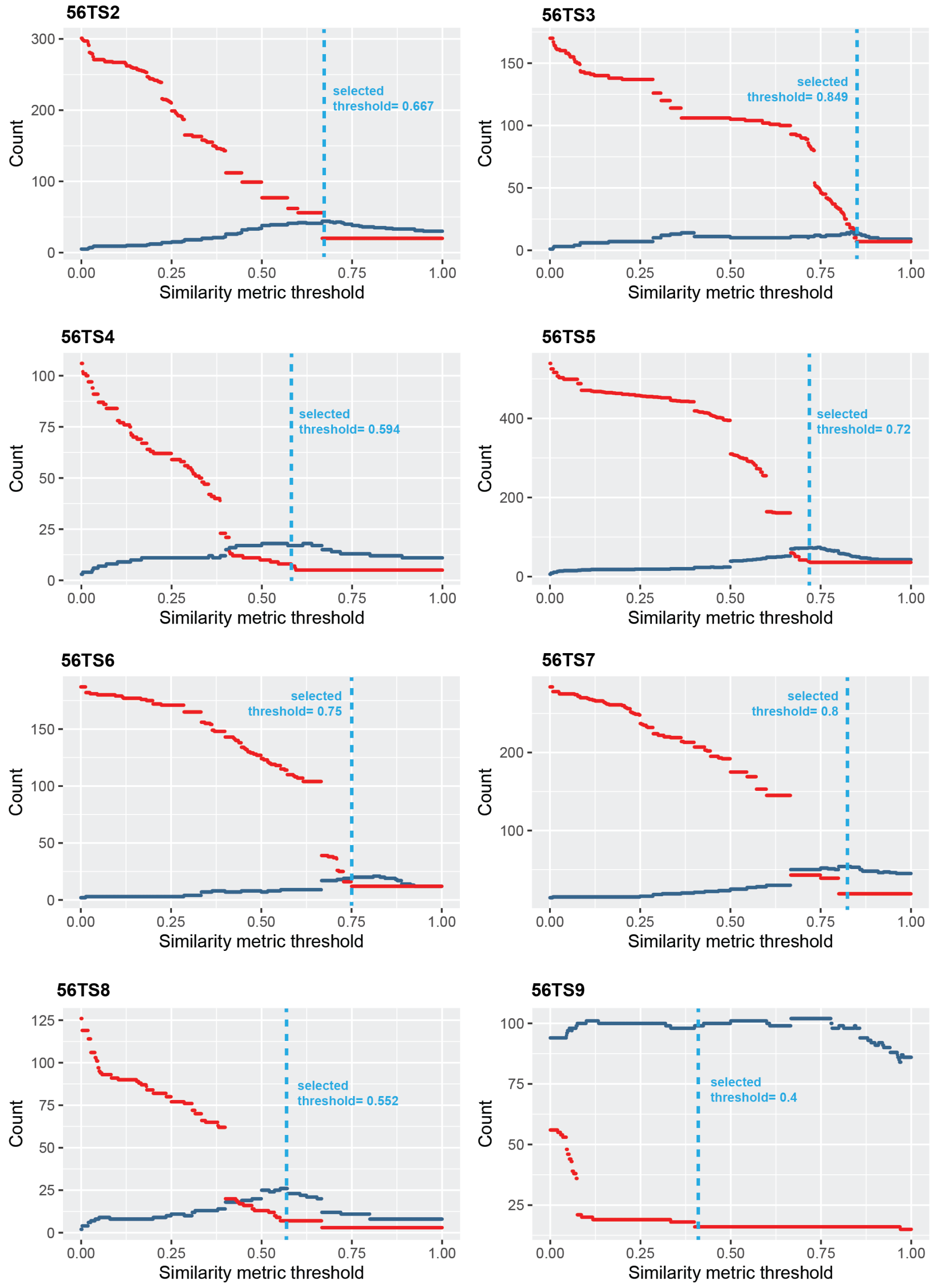


**fig. S4. TrackerSeq clone assignment statistics**

TrackerSeq clone assignment statistics according to edge weight filtering thresholds for each TrackerSeq experimental replicate generated in the study. Red indicates the size of the largest multicell clone detected in the dataset if the network of connected components is filtered for edge weights greater than the threshold as indicated on the x-axis. Blue indicates the number of unique multicell clones based on assignment using networks filtered for edge weights greater than the threshold value shown. The dashed line indicates the threshold selected for each dataset.

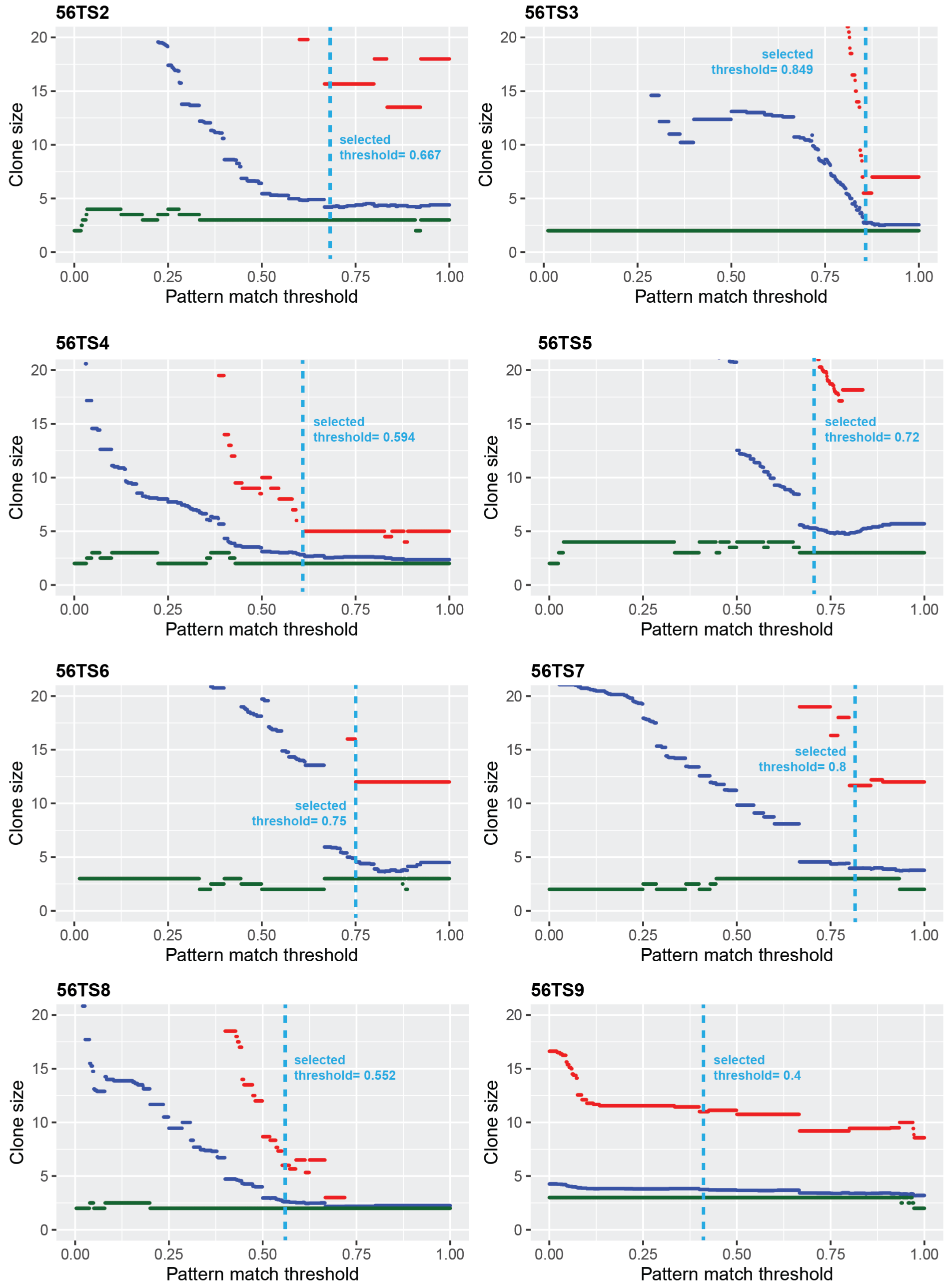


**fig. S5. TrackerSeq clone size statistics**

TrackerSeq clone size statistics according to edge weight filtering thresholds for each TrackerSeq experimental replicate generated in the study. Red indicates the mean size of the largest 10% of multicell clones detected in the dataset if the network of connected components is filtered for edge weights greater than the threshold as indicated on the x-axis. Blue indicates mean and green the median size of all multicell clones detected using the network at each threshold level.

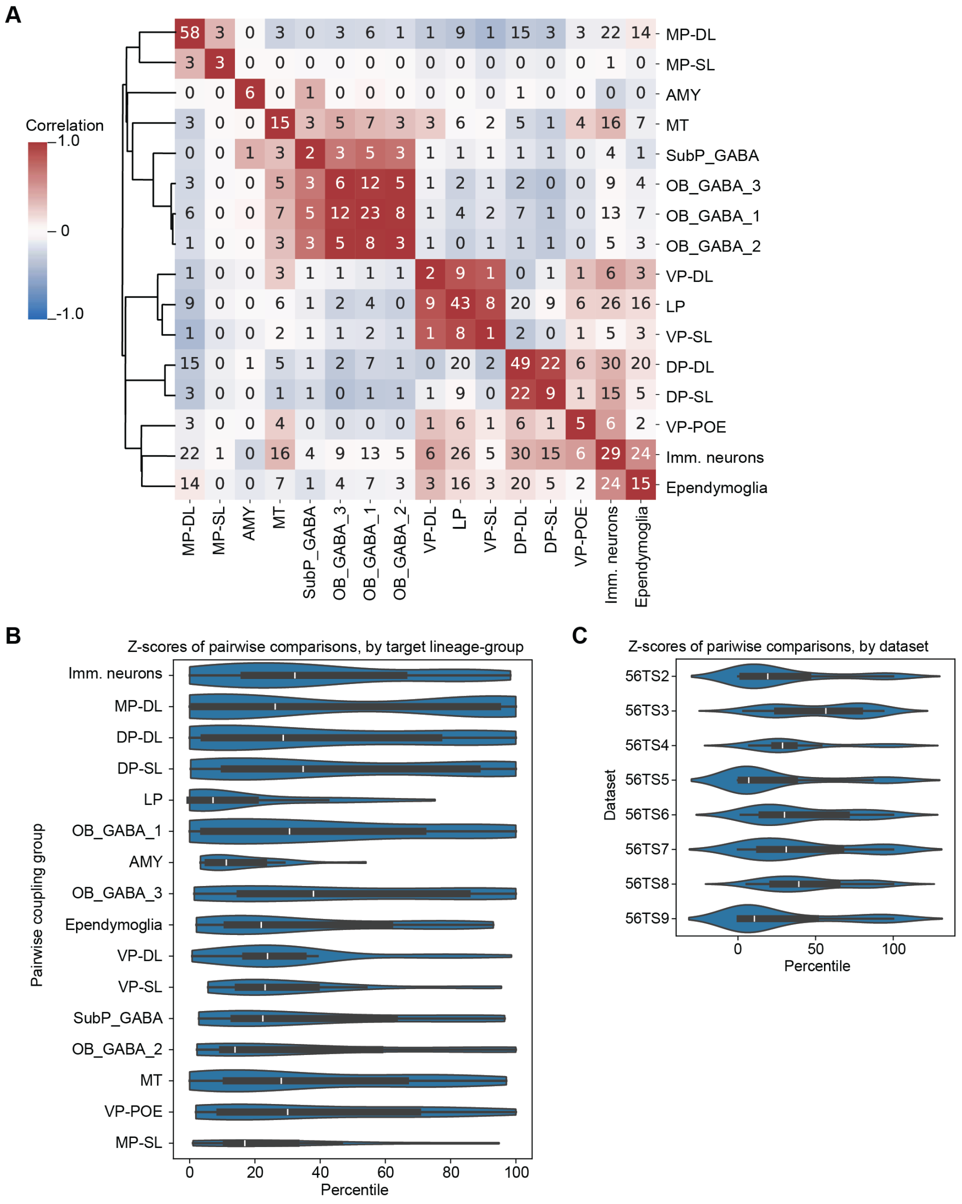


**fig. S6. TrackerSeq lineage correlation**

(**A**) Heatmap of pairwise lineage correlations based on lineage coupling z-score analysis. Multiple cell groups where pairwise correlation scores cluster together are more likely to appear together in the same clones. After z-scores are generated as in figure 2F, correlations are computed for each z-score pair, with high correlation indicating that cells are more likely to appear in clones together, and low correlation indicating that cells are less likely to appear in clones together; numbers indicate count of clonal intersections for each pair of cell-groups. (**B**) Distribution of z-scores by coupled cell type group, generated in lineage coupling analysis conducted independently from 8 experimental replicates. To compare z-scores from experimental replicates with different observation depths, z-scores were converted to percentiles. Each row indicates the distribution of lineage coupling scores between all cell type groups and the pairwise target group. Variance of these distributions was greater than the variance observed by dataset, as shown in C. (**C**) Distribution of z-scores by coupled experimental replicates, generated in lineage coupling analysis conducted independently from 8 experimental replicates. The variance of z-scores was significantly lower across datasets than across pairwise target groups, as shown in B.

Abbreviations: AMY, amygdala; DP-DL, dorsal pallium deep-layer neurons; DP-SL, dorsal pallium superficial-layer neurons; Imm. neurons, immature neurons; LP, lateral pallium; MP-DL, medial pallium deep-layer neurons; MP-SL, medial pallium superficial-layer neurons; MT, mitral and tufted cells of the olfactory bulb; OB_GABA_1-3, olfactory bulb GABAergic neurons; SubP_GABA, subpallium GABAergic neurons; VP-DL, ventral pallium deep-layer neurons; VP-POE, ventral pallium post-olfactory eminence; VP-SL, ventral pallium superficial-layer neurons

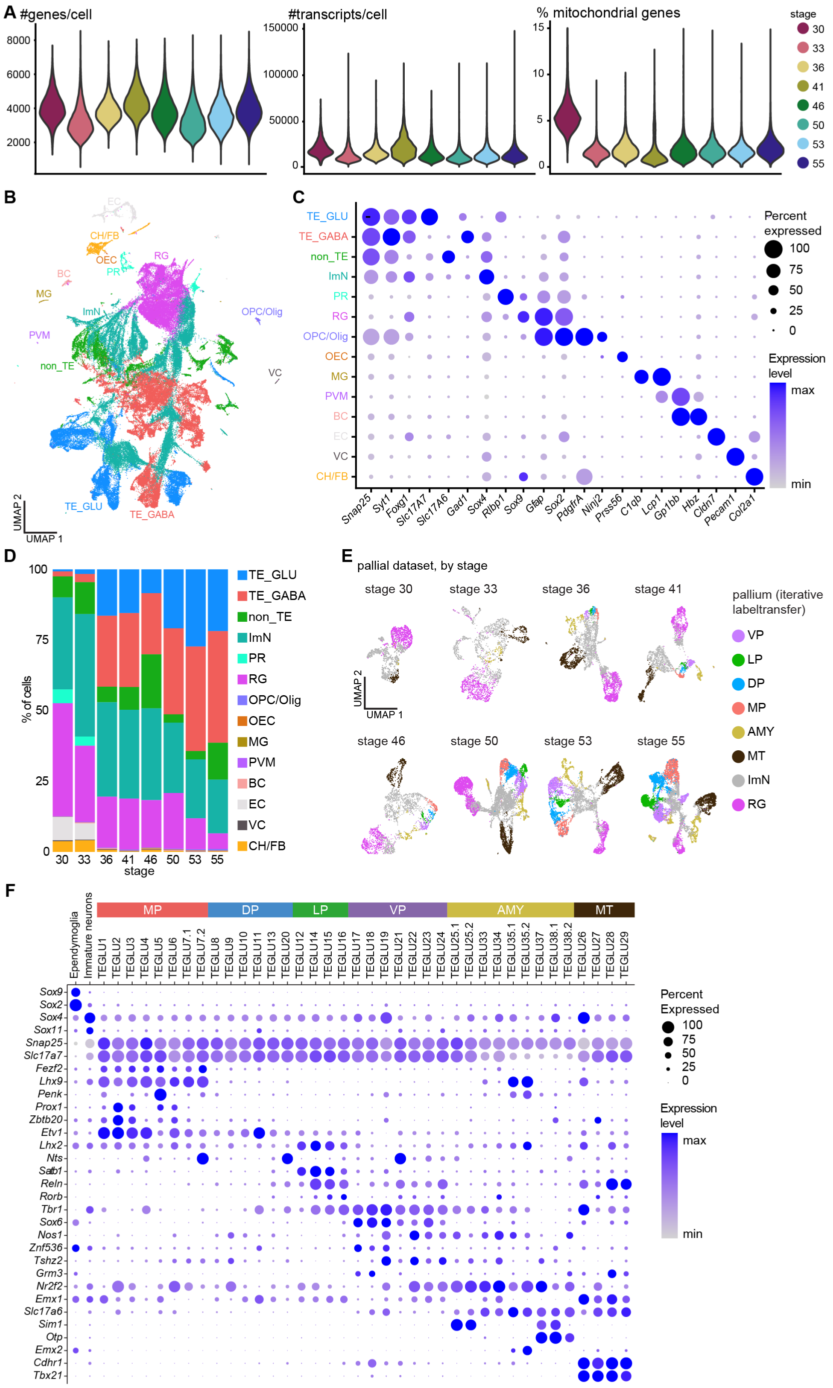


**fig. S7. Quality control of the developmental dataset.**

(**A**) Violin plots showing the number of genes, number of transcripts and percent mitochondrial genes per cell, grouped by stage. (**B**) UMAP plot of the developmental dataset, colored by cell class. (**C**) Dotplot showing the expression of marker genes used to annotate the developmental dataset in (B). (**D**) Barplot showing the percentage of cells in each class, depending on developmental stage. (**E**) UMAP plots of the pallial dataset for each sampling stage, colored by labeltransfer. (**F**) Dotplot of clusters annotated using LabelTransfer (see Methods), showing the expression of key marker genes defining distinct pallial regions, based on our previously published dataset.[46,47]

Abbreviations: AMY, amygdala; BC, blood cells; CH/FB, chondrocytes/fibroblasts; DL, deep-layer neurons; DP, dorsal pallium; EC, epithelial cells; ImN, immature neurons; LP, lateral pallium; MG, microglia; MP, medial pallium; MT, mitral and tufted cells; non_TE, non telencephalic neurons; OEC, olfactory ensheathing cells; OPC/Olig, oligodendrocyte precursor cells/oligodendrocytes; PVM, perivascular macrophages; PR, photoreceptor cells; RG, radial glia; SL, superficial-layer neurons; TE_GABA, telencephalic GABAergic neurons; TE_GLU, telencephalic glutamatergic neurons; VC, vascular cells; VP, ventral pallium

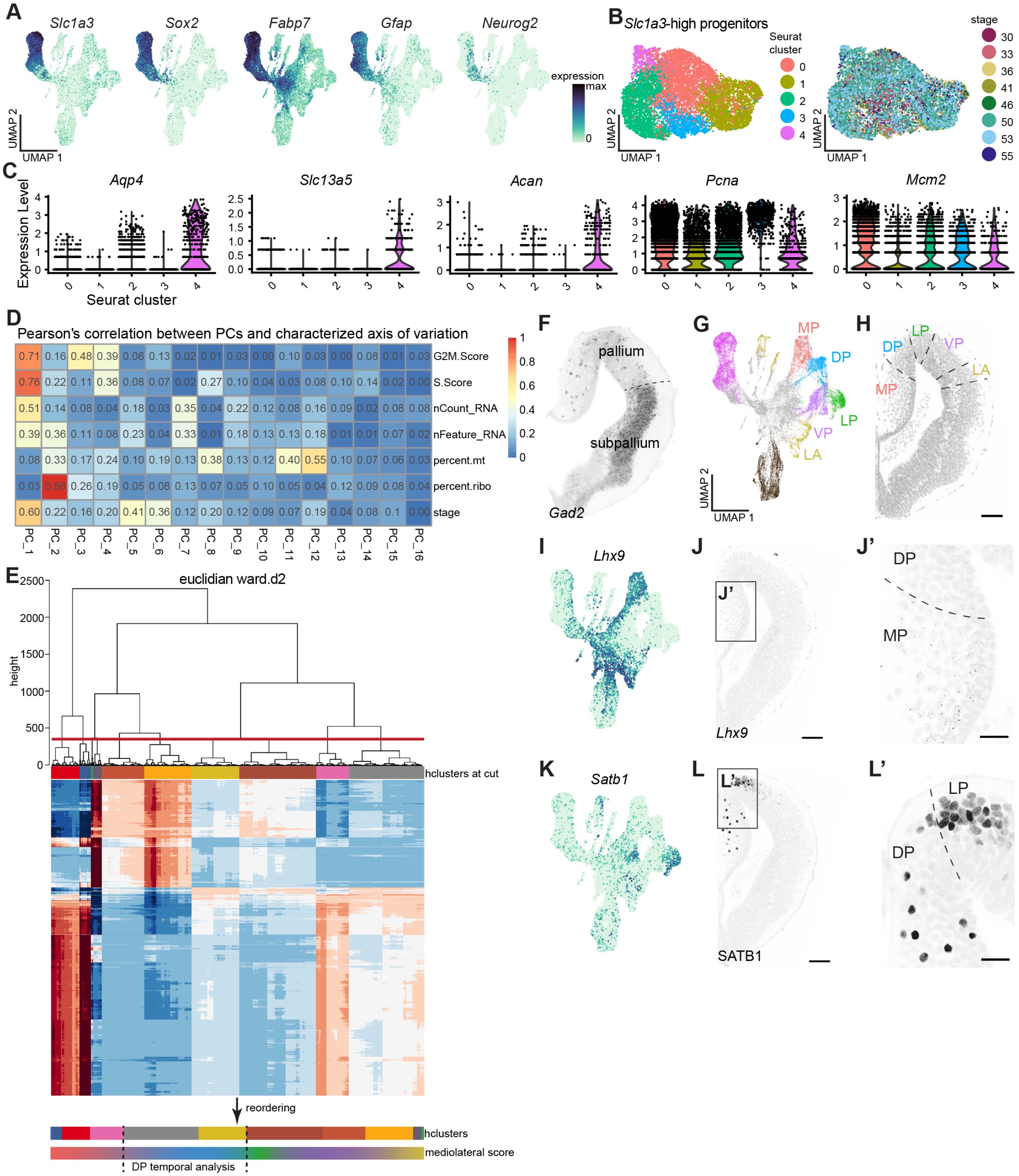


**fig. S8. Spatial heterogeneity of pallial progenitors.**

(**A**) UMAP plots of the pallium, colored by the expression of *Slc1a3*, *Sox2*, *Fabp7*, *Gfap* (progenitors), and *Neurog2* (neurogenic progenitors). (**B**) UMAP plots of the subsetted progenitor dataset colored by Seurat cluster (left) and developmental stage (right). (**C**) Violin plots of the subsetted progenitor dataset showing the expression of astrocyte markers *Aqp4*,

*Slc13a5*, *Acan* and proliferation markers *Pcna* and *Mcm2*. (**D**) Heatmap showing Pearson's correlation between PCs and known variables before regression. (**E**) Euclidian ward.d2 tree and heatmap of the final progenitor dataset, showing 10 clusters at the cut (red line). Clusters were re-ordered according to the Wnt to anti-Wnt gradient (Figure 5E), to identify DP progenitors for temporal analysis. (**F-L**) Visualization of the pallial territories in the stage 50 telencephalon using markers for differentiated GABAergic neurons (*Gad2*), medium pallium neurons (*Lhx9*) and lateral pallium neurons (SATB1).

Abbreviations: DP, dorsal pallium; LA, lateral amygdala; LP, lateral pallium; MP, medial pallium; VP, ventral pallium

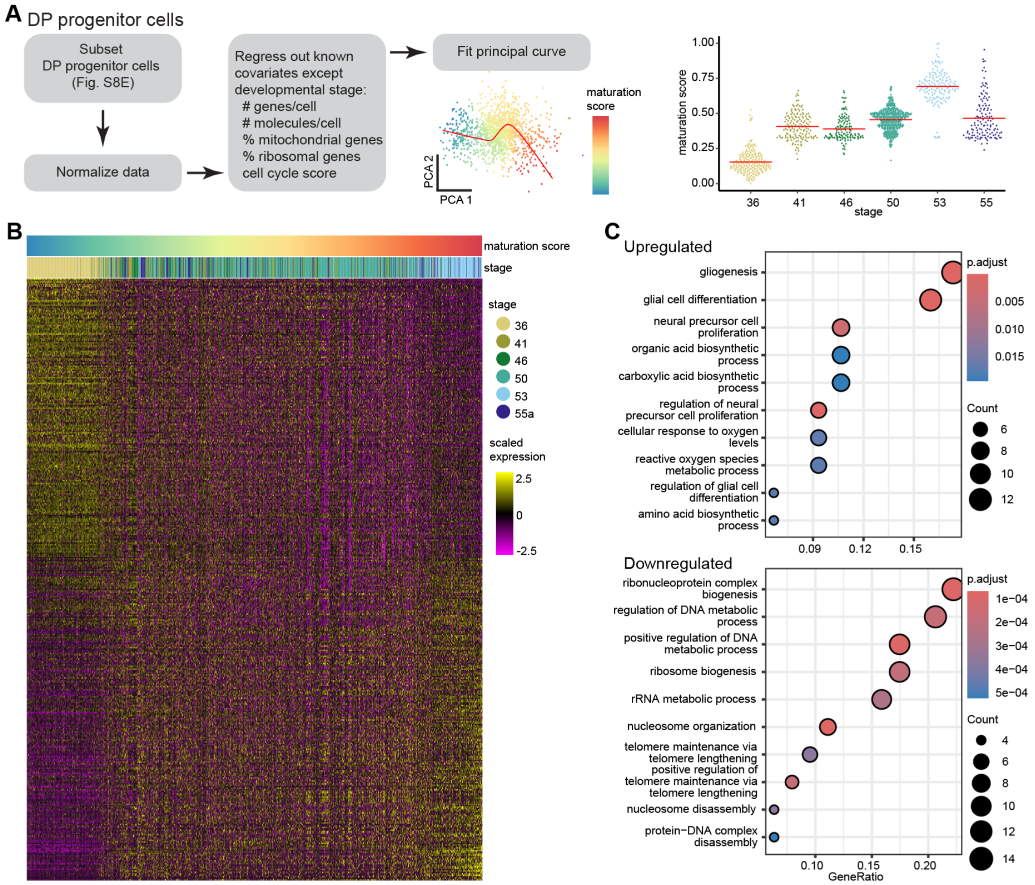


**fig. S9. Temporal heterogeneity of dorsal pallial progenitors.**

(**A**) Schematic representation of the temporal analysis for Figure 4F-I. (**B**). Heatmap showing differentially expressed genes along the maturation axis in the salamander DP. (**C**) GO analysis of shared salamander-mouse differentially expressed genes.

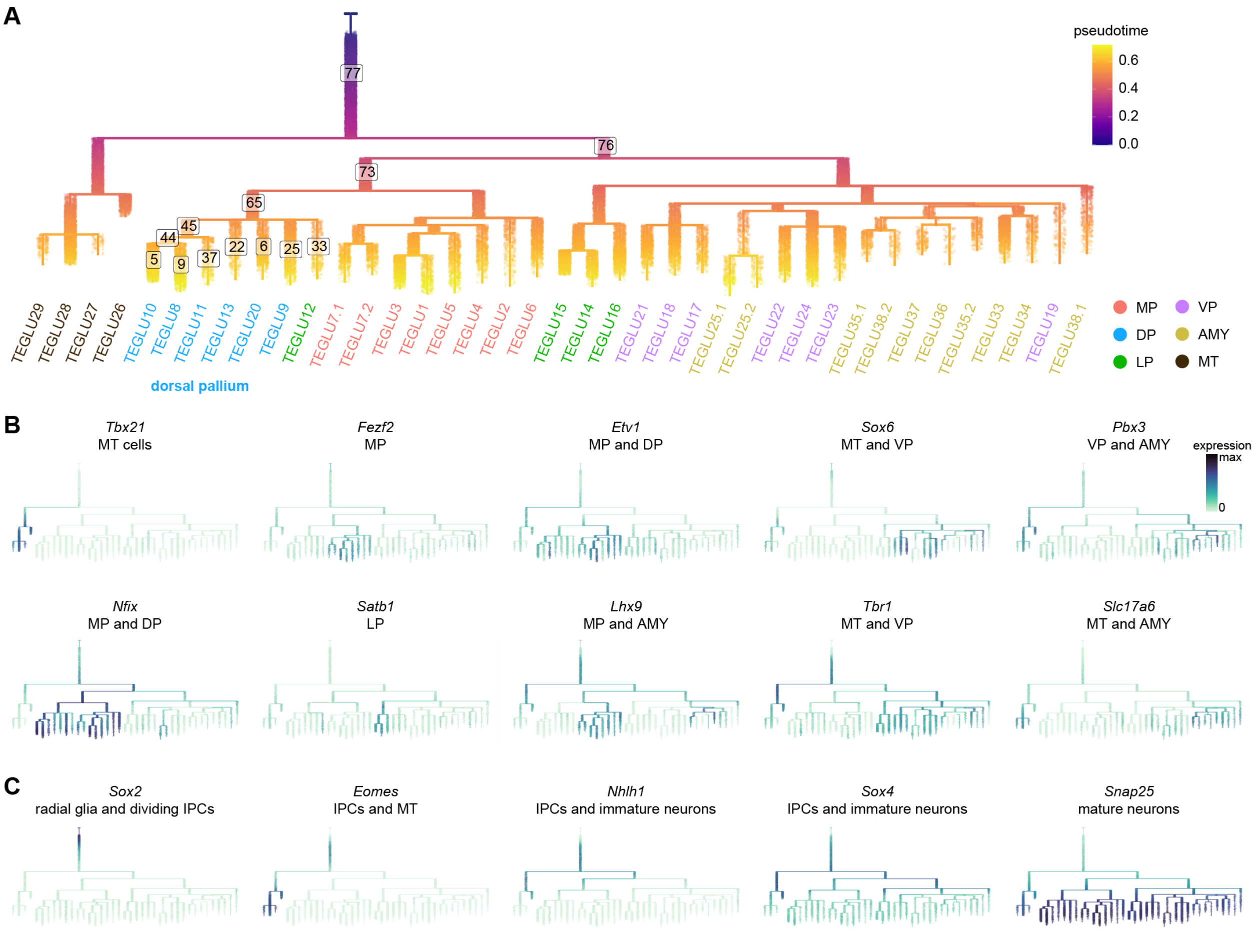


**fig. S10. URD analysis of salamander pallial development.**
(**A**) URD tree color-coded by pseudotime score. Segments used for further analysis of dorsal pallium trajectories are highlighted. (**B**) URD trees color-coded by the expression of marker genes of the different pallial domains. (**C**) URD trees color-coded by the expression of marker genes along the neurogenic trajectory.

Abbreviations: AMY, amygdala; DP, dorsal pallium; IPCs, intermediate progenitor cells; LP, lateral pallium; MP, medial pallium; MT, mitral and tufted cells of the olfactory bulb; TEGLU, telencephalic glutamatergic neurons; VP, ventral pallium

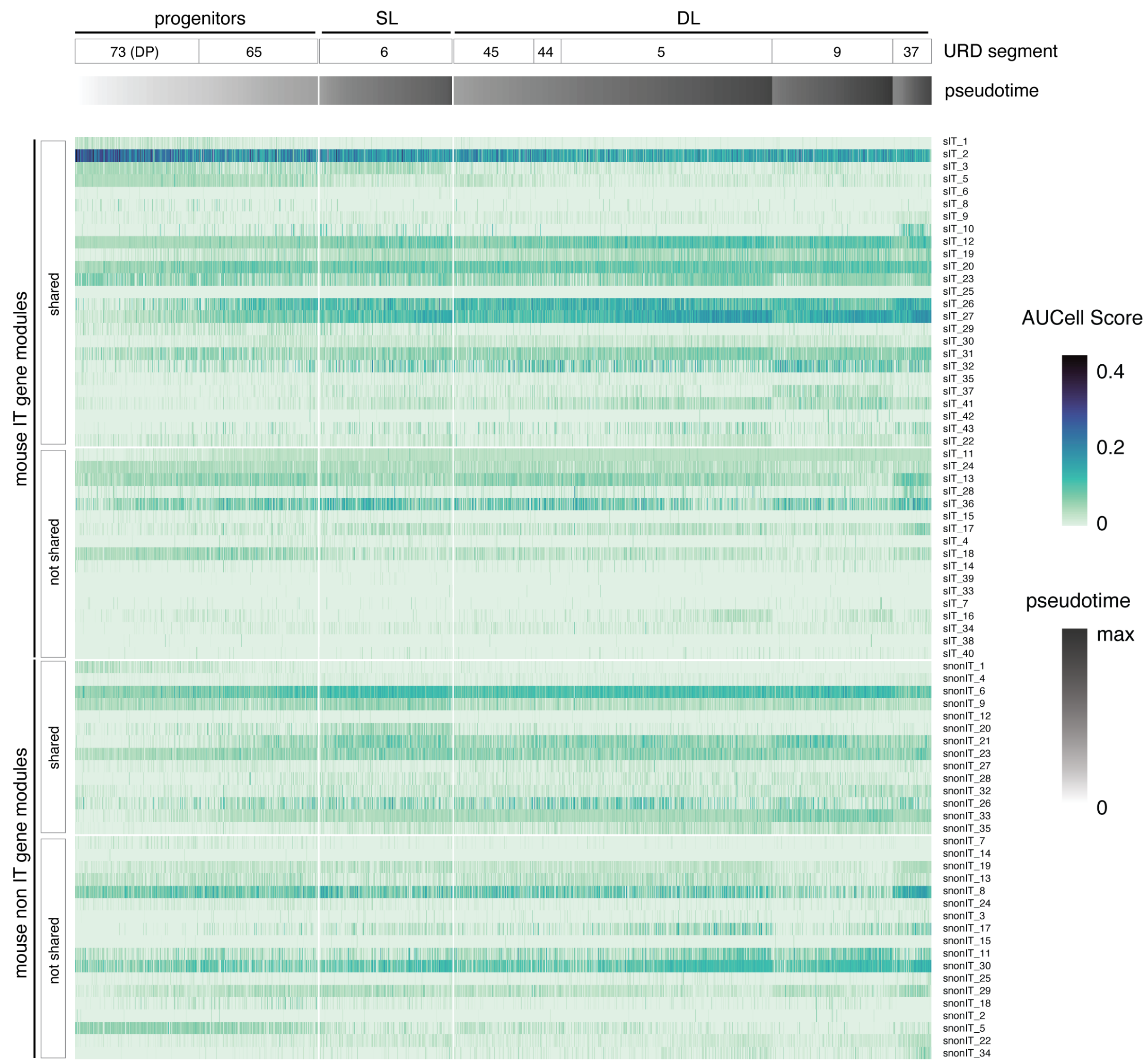


**fig. S11. Expression of developing mouse neocortex gene modules in salamander dorsal pallium.**

AUCell enrichment scores for gene modules identified during mouse neocortical development by Gao et al.[74] in cells across salamander dorsal pallium URD segments (fig. S10A). In mouse, these gene modules distinguish developmental trajectories of IT and non-IT neuronal classes. "Shared" refers to modules expressed in every cell type of the class, and "non shared" refers to cell-type specific modules. In salamanders, these modules do not have cell-type specific expression in SL and DL cells.

Abbreviations: DL, deep-layer; DP, dorsal pallium; IT, intratelencephalic; SL, superficial-layer

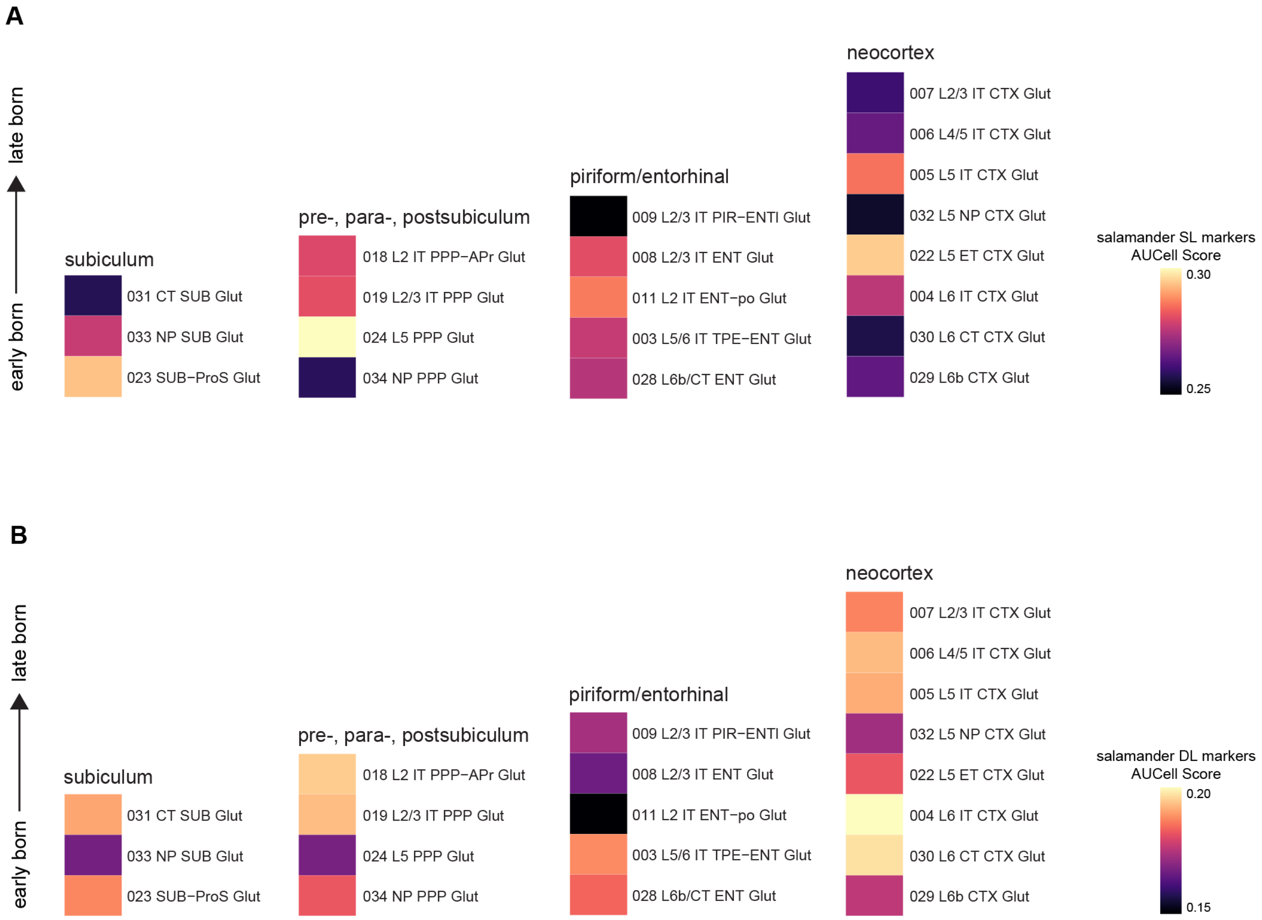


**fig. S12. AUCell enrichment scores for salamander SL and DL markers in mouse cortical cell types.**

(**A**) Salamander SL markers are enriched in mouse extratelencephalic projection neurons of the neocortex, subiculum, postsubiculum and retrosplenial cortex, while (**B**) salamander DL markers are generally enriched in mouse intratelencephalic-projecting neurons. Mouse data from Yao et al.[76].

Abbreviations: APr, area prostriata; CT, corticothalamic; CTX, cortex; ENT, entorhinal; ET, extratelencephalic; IT, intratelencephalic; L, layer; NP, near-projecting; PIR, piriform; PPP, pre-, para-, postsubiculum; ProS, prosubiculum; SUB, subiculum; TPE, transition piriform entorhinal area

*Supplementary tables*

**table S1. Number of shared clones contributing to each clone for each pairwise group (separate file)**

**table S2. Number of cells contributing to shared clones for each pairwise group (separate file)**

**table S3. Temporally differentially expressed genes in the salamander dorsal pallium (separate file)**

**table S4. Conserved genes showing differential expression along the temporal axis in mouse neocortex and salamander dorsal pallium (separate file)**
